# An atlas of epithelial cell states and plasticity in lung adenocarcinoma

**DOI:** 10.1101/2022.05.13.491635

**Authors:** Guangchun Han, Ansam Sinjab, Zahraa Rahal, Anne M. Lynch, Warapen Treekitkarnmongkol, Yuejiang Liu, Alejandra G. Serrano, Jiping Feng, Ke Liang, Khaja Khan, Wei Lu, Sharia D. Hernandez, Yunhe Liu, Xuanye Cao, Enyu Dai, Guangsheng Pei, Jian Hu, Camille Abaya, Lorena I. Gomez-Bolanos, Fuduan Peng, Minyue Chen, Edwin R. Parra, Tina Cascone, Boris Sepesi, Seyed Javad Moghaddam, Paul Scheet, Marcelo V. Negrao, John V. Heymach, Mingyao Li, Steven M. Dubinett, Christopher S. Stevenson, Avrum E. Spira, Junya Fujimoto, Luisa M. Solis, Ignacio I. Wistuba, Jichao Chen, Linghua Wang, Humam Kadara

## Abstract

Understanding cellular processes underlying early lung adenocarcinoma (LUAD) development is needed to devise intervention strategies^1^. Here we studied 246,102 single epithelial cells from 16 early-stage LUADs and 47 matched normal lung samples. Epithelial cells comprised diverse normal and cancer cell states, and diversity among cancer cells was strongly linked to LUAD-specific oncogenic drivers. KRAS mutant cancer cells showed distinct transcriptional features, reduced differentiation and low levels of aneuploidy. Non-malignant areas surrounding human LUAD samples were enriched with alveolar intermediate cells that displayed elevated KRT8 expression (termed KRT8+ alveolar intermediate cells (KACs) here), reduced differentiation, increased plasticity and driver KRAS mutations. Expression profiles of KACs were enriched in lung precancer cells and in LUAD cells and signified poor survival. In mice exposed to tobacco carcinogen, KACs emerged before lung tumours and persisted for months after cessation of carcinogen exposure. Moreover, they acquired Kras mutations and conveyed sensitivity to targeted KRAS inhibition in KAC-enriched organoids derived from alveolar type 2 (AT2) cells. Last, lineage-labelling of AT2 cells or KRT8+ cells following carcinogen exposure showed that KACs are possible intermediates in AT2-to-tumour cell transformation.

## Main

Lung adenocarcinomas (LUAD) are increasingly being detected at earlier pathological stages due to enhanced screening^2–4^. Yet, patient prognosis remains moderate to poor thus warranting improved early treatment strategies. Decoding the earliest events driving LUADs can inform of ideal targets for its interception. Previous work showed that smoking leads to pervasive molecular (e.g., *KRAS* mutations) and immune changes that are shared between LUADs and their adjacent normal-appearing ecosystems and that are strongly associated with development of lung premalignant lesions (PMLs) and LUAD^1,5–12^. Yet, most of these reports were based on bulk approaches and were designed to focus on tumour and distant normal sites in the lung, thereby discounting the cellular and transcriptional phenotypes of the expanded LUAD landscapes. Furthermore, while many lung single-cell RNA-sequencing (scRNA-seq) studies have decoded immune and stromal states^13,14^, little insight is drawn to epithelial cells given their paucity (∼4%) when performing single-cell analyses without enrichment of the epithelial compartment. Therefore, little is known of the identities of specific epithelial subsets or how they promote a field of injury, trigger progression of normal lung (NL) to PML and inspire LUAD pathogenesis. Understanding cell type-specific changes at the root of LUAD inception will help identify actionable targets and strategies for prevention of this morbid disease. Here, our efforts were focused on in-depth single-cell interrogation of malignant and normal epithelial cells from both early-stage LUADs as well as carcinogenesis and lineage tracing mouse models that recapitulate the disease, with attention to how specific populations evolve to give rise to malignant tumours.

### Epithelial transcriptional landscape

Our study design converges in-depth scRNA-seq of early-stage LUAD clinical specimens as well as cross-species analysis and lineage tracing in a human-relevant model of LUAD development following exposure to tobacco carcinogen (**Fig. 1a**). We studied by scRNA-seq EPCAM-enriched epithelial cell subsets from early-stage LUADs of 16 patients and 47 paired NL samples spanning a topographic continuum from the LUADs, that is, tumour-adjacent, -intermediate, and -distant locations^15^ (**Fig. 1a, Supplementary Fig. 1,** and **Supplementary Tables 1, 2**). We also collected from the same regions, tumour and normal tissue sets for whole exome sequencing (WES) profiling and for high-resolution spatial transcriptomics (ST) and protein analyses (**Fig. 1a**).

**Fig. 1.**
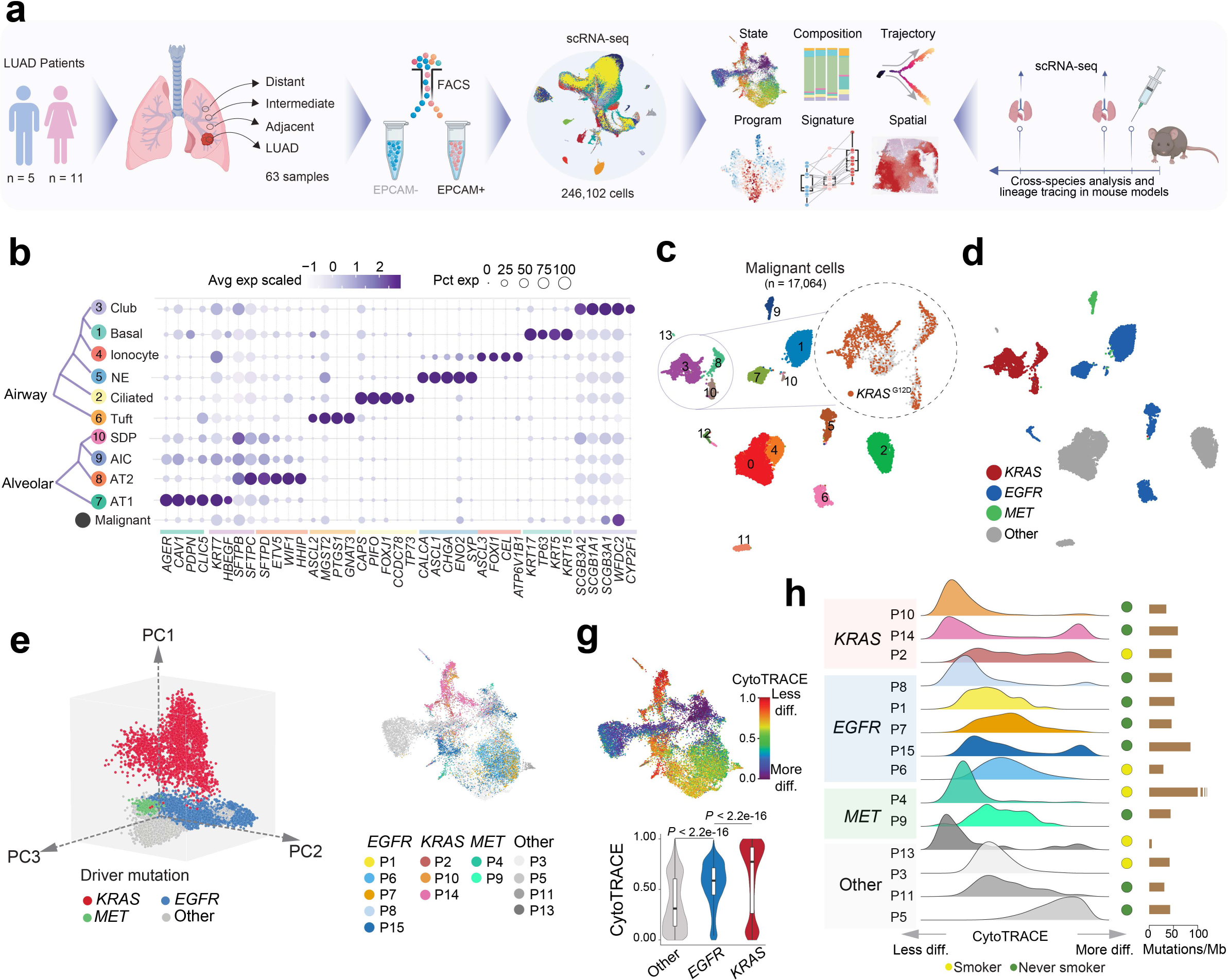
Transcriptional landscape of lung epithelial and malignant cells in early-stage LUAD. **a,** Schematic overview of experimental design and analysis workflow. State: cellular transcriptional state. Composition: composition of cell subsets. Program: transcriptional programs in malignant cells. Spatial: *in situ* spatial transcriptome and protein analyses. Created with BioRender.com. **b,** Proportions and average expression levels) of selected marker genes for ten normal epithelial and one malignant cell subset. **c,** Unsupervised clustering of 17,064 malignant cells coloured by cluster ID. top right inlet shows malignant cells coloured by *KRAS*^G12D^ mutation status identified by scRNA-seq. **d,** Uniform Manifold Approximation and Projection (UMAP) of malignant cells shown in **c** and coloured by driver mutations identified in each tumour sample using WES. **e,** Principal component analysis (PCA) plot of malignant cells coloured by driver mutations identified in each tumour sample by WES. **f,** UMAP plots of malignant cells coloured by patient ID and grouped by driver mutation status. **g,** UMAP of malignant cells by differentiation state inferred by CytoTRACE (top). Comparison of CytoTRACE scores between malignant cells from samples with different driver mutations (bottom), box, median ± interquartile range; whiskers, 1.5× interquartile range; centre line: median. *n* cells in each box-and-whisker (left to right): 9,135; 5,457; 2,472. *P* - values were calculated using a two-sided Wilcoxon rank-sum test with a Benjamini–Hochberg correction. **h**, Per sample distribution of malignant cell CytoTRACE scores.

Following quality control, 246,102 epithelial cells were retained (**Supplementary Fig. 2** and **Supplementary Table 2**). Malignant cells (n = 17,064) were distinguished from otherwise non-malignant “normal” cells (n = 229,038) by integrating information from inferred copy number variations (inferCNV^16^), clustering distribution, lineage-specific gene expression, and the presence of reads carrying *KRAS*^G12D^ somatic mutations (**Fig. 1b, c** and **Supplementary Fig. 3**). Analysis of non-malignant clusters identified two major lineages, namely alveolar and airway, as well as a small subset of proliferative cells (**Extended Data Fig. 1a** and **Supplementary Table 2**). Airway cells (n = 40,607) included basal (*KRT17^+^*), ciliated (*FOXJ1^+^*), and club and secretory (*SCGB1A1^+^*) populations, as well as rare types such as ionocytes (*ASCL3^+^*), neuroendocrine (*ASCL1^+^*), and tuft cells (*GNAT3^+^*) (**Extended Data Fig. 1a** and **Supplementary Table 3**). Alveolar cells (n = 187,768) consisted of alveolar type 1 (AT1; *AGER1^+^*, *ETV5^+^*), alveolar type 2 (AT2; *SFTPB^+^, SFTPC^+^*), *SCGB1A1/SFTPC* dual positive cells (SDP), as well as a cluster of alveolar intermediate cells (AICs) that was closely tucked between AT1 and AT2 clusters and that shared gene expression features with both major alveolar cell types (**Fig. 1b** and **Extended Data Fig. 1a, b**).

Malignant cells showed low/no expression of lineage-specific markers and, overall, diminished lineage identity (**Fig. 1b**, **bottom**). Malignant cells formed 14 clusters (**Fig. 1c**) that were for the most part patient-specific (**Extended Data Fig. 1c, left**) signifying strong interpatient heterogeneity. Overall, malignant cells showed high levels of aneuploidy (**Extended Data Fig. 1c, middle**). We did not detect any unique clustering pattern with respect to smoking status (**Extended Data Fig. 1d**). Annotation based on genomic profiling (by WES) showed that malignant cells from 3 patients with *KRAS*-mutant (KM-) LUADs (P2, P10, P14) clustered closely together and in comparison to malignant cells from other LUADs which showed a more dispersed clustering pattern (**Fig. 1d, Extended Data Fig. 1c, e** and **Supplementary Table. 1**). scRNA-seq analysis confirmed the presence of CNVs and *KRAS*^G12D^ mutations in patient-specific tumour clusters as well as absence of *KRAS*^G12D^ in *KRAS* wild type (KW-) LUADs (**Extended Data Fig. 1c**).

### LUAD malignant transcriptional programs

Malignant cells from KM-LUADs clustered together and distinctively from those of *EGFR*- mutant (EM-) or *MET*-mutant (MM-) LUADs (**Fig. 1e**). KM-LUADs showed more transcriptomic similarity (i.e., shorter Bhattacharyya distances) at both sample and cell levels (**Extended Data Fig. 1f, left** and **right,** respectively) compared to other LUADs (*P* < 2.2 × 10^-16^). Distances between KM-LUAD samples (KM-KM) were significantly smaller compared to those between EM-LUADs (EM-EM; *P* = 0.02) or other LUADs (Other-Other; *P* = 0.03; **Extended Data Fig. 1f, left**). Clustering of malignant cells, following adjustment for patient-specific effects, showed that cluster 5 was enriched with cells from KM-LUADs (P2, P10 and P14; **Extended Data Fig. 1g**). Most of the *KRAS*-mutant malignant cells clustered separately from other cells, indicating unique transcriptional programs in *KRAS*-mutant cells (**Fig. 1f**). In line with previous reports^15,17^, malignant cells from KM-LUADs were chromosomally more stable in contrast to those from EM-LUADs (*P* < 2.2 × 10^-16^; **Extended Data Fig. 1h, left**). CNV events were significantly more prominent in malignant cells from smoker relative to never-smoker patients (*P* < 2.2 × 10^-16^; **Extended Data Fig. 1h, right**). Differentiation states of malignant cells exhibited high inter-patient heterogeneity, whereby, irrespective of tumour mutation load, KM-LUAD cells were the least differentiated as indicated by their highest CytoTRACE^18^ scores, followed by EM-LUADs (*P* < 0.001; **Fig. 1g, h** and **Supplementary Table 4**). There was intra-tumour heterogeneity (ITH) in differentiation states (e.g. P2, P9, P14, P15), whereby malignant cells from 7 out of the 14 patients with detectable malignant cells exhibited a broad distribution of CytoTRACE scores, with KM-LUADs showing a trend for higher variability in differentiation (greater Wasserstein distances) relative to EM- or other LUADs (**Fig. 1h** and **Extended Data Fig. 1i**).

Clustering of malignant cells (Meta_C1 to C5) based on levels of 23 recurrent meta-programs (MPs)^19^ showed that Meta_C1 comprised cells mostly from KM-LUADs (92%) and displayed highest expression of gene modules associated with *KRAS*^G12D^ present in pancreatic ductal adenocarcinoma (MP30)^19^, epithelial to mesenchymal transition (EMT-III; MP14) and epithelial senescence (MP19), and, conversely, lowest levels of alveolar MP (MP31) (**Extended Data Fig. 2a-c, Supplementary Table 5**). Notably, malignant cells from patients P2, P10, and P14 with KM-LUADs showed significantly higher expression of MP30 relative to those from patients with KW-LUADs (*P* < 2.2 x 10^-16^; **Extended Data Fig. 2d**). Malignant cell states also exhibited ITH in KM-LUAD (e.g., P14; **Extended Data Fig. 2e**). A subset of *KRAS*^G12D^ cells showed activation of MP30 and there were diverse activation patterns for other MPs (e.g., cell respiration) across the mutant cells (**Extended Data Fig. 2e middle, f**). Overall, malignant cells carrying *KRAS*^G12D^ showed reduced differentiation (**Extended Data Fig. 2e, right**) which was concordant with loss of alveolar differentiation (MP31) in KM-LUADs (**Extended Data Fig. 2a, b**). P14 malignant cell clusters exhibited different levels of CNVs^15^, whereby a cluster enriched with *KRAS*^G12D^ cells harboured relatively late CNV events (e.g. Chr 1p loss, Chr 8 and Chr 12 gains) and reduced alveolar signature scores, in line with attenuated differentiation (**Extended Data Fig. 2g, h**). A KRAS signature was derived based on unique expression features of *KRAS*-mutant malignant cells from our cohort (i.e., specific to cluster 5; **Extended Data Fig. 1g**), and found to strongly and significantly correlate with MP30 signature (R = 0.92, *P* < 2.2 × 10^-16^, **Extended Data Fig. 2i** and **Supplementary Table 6**). KM-LUADs from TCGA cohort and with relatively high expression of our KRAS signature were enriched with activated *KRAS* MP30 as well as other MPs we had found to be increased in Meta_C1 (**Extended Data Fig. 2j**). KW-LUADs in TCGA with relatively higher expression of the KRAS signature displayed significantly lower overall survival (OS; *P =* 0.02; **Extended Data Fig. 2k**). A similar trend was observed when analysing *KRAS*^G12D^-mutant LUADs alone despite the small cohort size (*P =* 0.3; **Extended Data Fig. 2k**). These data highlight extensive transcriptomic heterogeneity between LUAD cells and transcriptional programs that are biologically- and possibly-clinically relevant to KM-LUAD.

### Alveolar intermediate cells in LUAD

In contrast to AT2 cells which were overall decreased in LUADs compared to multi-region NLs (*P* = 0.002), AICs showed the opposite pattern (*P* = 0.02; **Extended Data Fig. 3a, b**). AT2 fractions were gradually reduced with increasing tumour proximity across multi-region NLs from 7 of the 16 patients with LUAD (*P* = 0.004; **Extended Data Fig. 3c, d**). No significant changes in fractions were found for other major lung epithelial cell types (**Extended Data Fig. 3e**). AICs were intermediary along the AT2-to-AT1 developmental and differentiation trajectories (**Fig. 2a** and **Extended Data Fig. 3f, g**), reminiscent of intermediary alveolar cells in cancer-free mice exposed to acute lung injury^20^. The proportion of least differentiated AICs in LUAD tissues was higher than that of their more differentiated counterparts (29% versus 11%, respectively, **Extended Data Fig. 3h**). Notably, AICs were inferred to transition to malignant cells, including *KRAS*-mutant cells that were more developmentally late relative to *EGFR*-mutant malignant cells (*P* < 2.2 × 10^-16^; **Fig. 2a** and **Extended Data Fig. 3f**). Further analysis of AICs identified a subpopulation with uniquely high expression of *KRT8* (**Fig. 2b**). These “*KRT8*^+^ Alveolar Intermediate Cells”, or KACs, had increased expression of *CDKN1A/2A, PLAUR,* and the tumour marker *CLDN4* (**Fig. 2b**, **Extended Data Fig. 3i** and **Supplementary Table 7**). KACs were also significantly less differentiated (*P* < 2.2 × 10^-16^; **Fig. 2c**) and more developmentally late (*P =* 1.2 × 10^-11^; **Extended Data Fig. 3j**). Notably, KACs transitioned to *KRAS*-mutant malignant cells in pseudotime, whereas other AICs were more closely associated with differentiation to AT1 cells (**Extended Data Fig. 3j**). Proportions of KACs among non-malignant epithelial cells was strongly and significantly increased in LUADs relative to multi-region NL tissues (*P =* 2.4 × 10^-4^; **Fig. 2d**), and were significantly higher in LUADs than AT1, AT2, or other AICs fractions (*P* < 2.2 × 10^-16^; **Fig. 2e**). Notably, tumour-associated KACs clustered farther away from AICs compared to NL-derived KACs (**Extended Data Fig. 3k**).

**Fig. 2.**
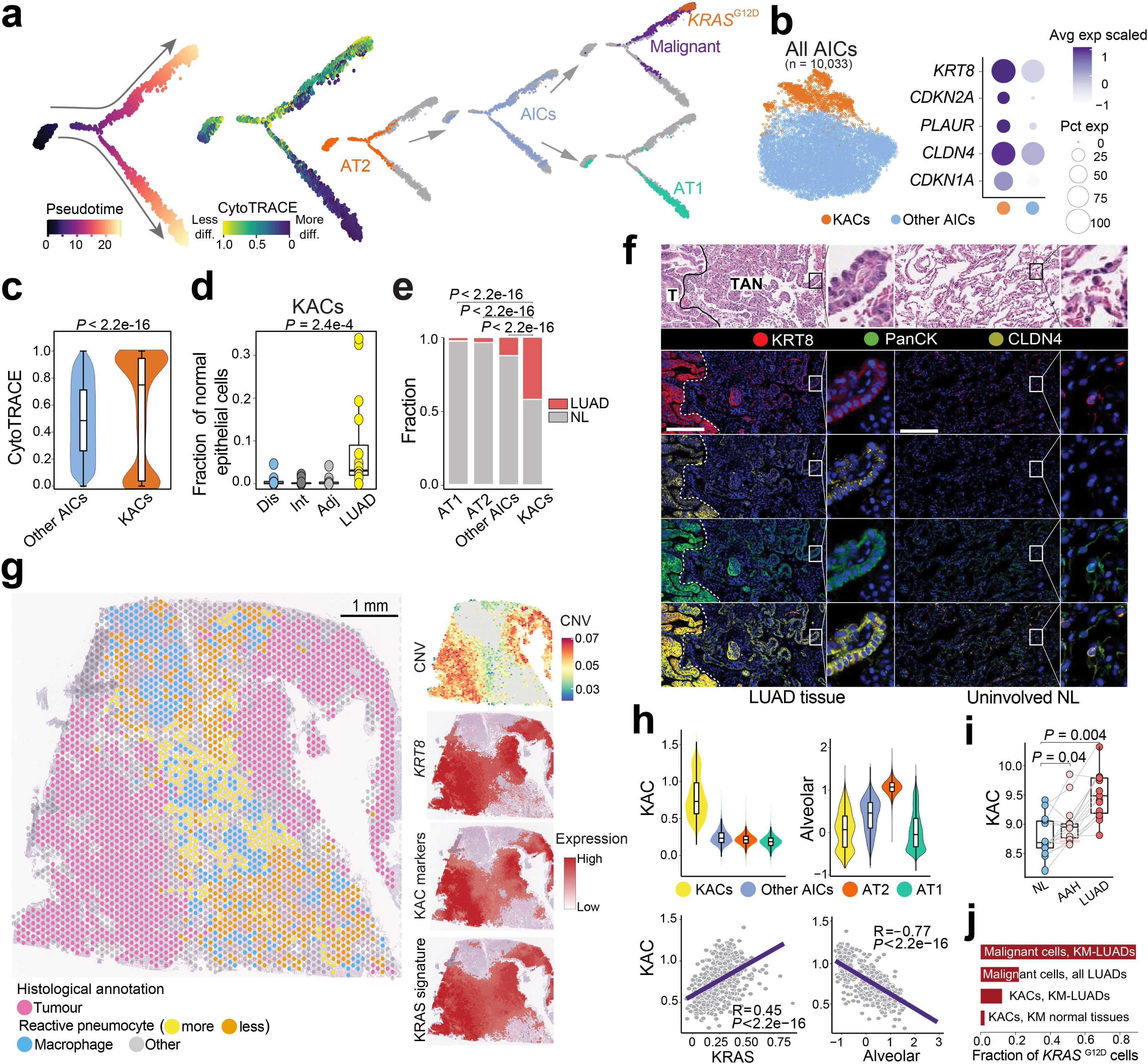
Identification and characterization of *KRT8*^+^ alveolar intermediate cells in human LUAD. **a,** Pseudotime analysis of alveolar and malignant cells. **b,** Sub-clustering analysis of AICs (left). Proportions and average expression levels of representative KAC marker genes (right). **c,** CytoTRACE score in KACs versus other AICs. Box-and-whisker definitions are similar to Fig. 1g with *n* cells (left to right): 8,591; 1,440. *P* - values were calculated using a two-sided Wilcoxon rank-sum test. **d,** Proportion of KACs among non-malignant epithelial cells. Box-and-whisker definitions are similar to **Fig. 1g** with *n* samples (left to right): 16; 15; 16; 16. *P* - value was calculated using Kruskal-Wallis test. **e,** Fraction of alveolar cell subsets coloured by sample type. *P* - values were calculated using two-sided Fisher’s exact tests with a Benjamini–Hochberg correction. **f,** Haematoxylin and Eosin (H&E) staining of LUAD tumour (T), TAN displaying reactive hyperplasia of AT2 cells, and uninvolved NL (top). Digital spatial profiling (DSP) shows KRT8, PanCK, CLDN4, Syto13 blue nuclear stain, and composite image (bottom). Magnification: 20X. Scale bar: 200 μm. Staining was repeated four times with similar results. Dashed white lines represent the margins separating tumours and TAN regions. **g,** ST analysis of P14 LUAD showing histologically annotated H&E-stained Visium slide (left) and spatial heatmaps depicting CNV score and scaled expression of *KRT8*, KAC markers (panel b), and KRAS signature. **h,** Expression and correlation analyses of KAC, KRAS and alveolar signatures. Box-and-whisker definitions are similar to **Fig. 1g** with *n* KACs = 1,440; Other AICs = 8,593; AT2 = 146,776; AT1 = 25,561. R: Spearman correlation coefficient. *P* - values were calculated with Spearman correlation test. **i,** KAC signature expression in premalignancy cohort (15 samples each). Box-and-whisker definitions are similar to **Fig. 1g**. *P* - values were calculated using two-sided Wilcoxon signed-rank test with a Benjamini–Hochberg correction. **j,** Fraction of *KRAS*^G12D^ cells in different subsets.

High-resolution, multiplex imaging analysis of KRT8, CLDN4, and pan-cytokeratin (PanCK) showed that KACs were not only enriched in tumour-adjacent normal regions (TANs) but were also found right next to malignant cells showing high expression of KRT8 and CLDN4 (**Fig. 2f** and **Extended Data Fig. 4a**). While KACs were also found in the uninvolved NLs, consistent with our scRNA-seq analysis, only in the TANs did they display features of “reactive” epithelial cells (**Fig. 2f** and **Extended Data Fig. 4a**). ST analysis of P14 tumour tissue demonstrated increased expression of *KRT8* in tumour regions (with high CNV scores), as well as in TAN regions that were histologically found to comprise highly reactive pneumocytes and that exhibited moderate/low CNV scores (**Fig. 2g**). Deconvolution showed that KACs were closer to tumour regions relative to alveolar cells (**Extended Data Fig. 4b**). ST analysis of a KAC-enriched region showed that KACs were indeed intermediary in the transition of alveolar parenchyma to tumour cells (**Extended Data Fig. 4b**). Tumour regions had markedly reduced expression of *NKX2-1* and alveolar signature (**Extended Data Fig. 4b**), in line with reduced alveolar differentiation in KM-LUADs (**Extended Data Fig. 2b**).

KAC markers (**Fig. 2b**) were high in tumour regions and in TANs with reactive pneumocytes as well as spatially overlapped with *KRAS* signature (**Fig. 2g**). Similar to KRAS but unlike AT1 and alveolar signatures, a KAC signature we derived was highest in KACs relative to AT1, AT2, or other AICs (**Fig. 2h**, **Extended Data Fig. 4c, d** and **Supplementary Table 8**). A signature pertinent to other AICs we derived was evidently lower in KACs relative to other AICs (**Extended Data Fig. 4e**). In KACs from all samples, KAC and KRAS signatures positively correlated together (R = 0.45; *P* < 2.2 × 10^-16^) and inversely so with their alveolar counterpart (R = -0.77; *P* < 2.2 × 10^-16^; **Fig. 2h**). In contrast, there was no correlation between “other AIC” and KRAS (R = 0.045; *P =* 3.2 × 10^-5^) nor alveolar (R = -0.11; *P* < 2.2 × 10^-16^) signatures (**Extended Data Fig. 4f, g**). The KAC signature was significantly higher in KACs and in malignant cells from KM-LUADs relative to those from EM-LUADs (*P* < 2.2 × 10^-16^; **Extended Data Fig. 4h**). In sharp contrast to “other AIC” and alveolar signatures, KAC signature was significantly enriched in TCGA LUADs compared to their matched uninvolved NLs (*P =* 1.9 × 10^-15^; **Extended Data Fig. 5a-c**). Of note, KAC signature was significantly and progressively increased along matched NL, premalignant atypical adenomatous hyperplasia (AAH) and invasive LUAD (**Fig. 2i**), whereas there was no such pattern for “other AIC” signature (**Extended Data Fig. 5d**). KAC signature was significantly higher in TCGA KM-LUADs relative to KW-LUADs (*P =* 0.002; **Extended Data Fig. 5e**). Also, KAC but not “other AIC” signature was significantly associated with reduced OS in two independent cohorts (TCGA, *P =* 0.005; PROSPECT, *P =* 0.04; **Extended Data Fig. 5f-i**). KAC signature was associated with shortened OS even after accounting for stage (FDR adjusted *q - value* = 0.034; **Extended Data Fig. 5j)**.

Despite exhibiting lower CNV scores compared to malignant cells, KACs exhibited moderately elevated CNV burdens relative to AT2, AT1, and other AICs (**Extended Data Fig. 6a, b**). *KRAS*^G12D^ was present in malignant cells with a variant allele frequency (VAF) of 78% in KM-LUADs (**Fig. 2j**, **Extended Data Fig. 6c** and **Supplementary Table 9**). KACs, but not AT2, AT1, or other AICs, harboured *KRAS*^G12D^ mutations (**Extended Data Fig. 6c, d**). *KRAS*^G12D^ KACs were exclusively found in tissues (primarily tumours) from KM-LUADs and, thus, *KRAS*^G12D^ VAF (10%) was higher in KACs from KM-LUADs compared to when examined in all LUADs (5%) or samples (3%) (**Fig. 2j** and **Extended Data Fig. 6c, d**). *KRAS*^G12D^ mutations were detected in KACs of NL tissues of patients with KM-LUAD (VAF 2%), and other *KRAS* variants (*KRAS*^G12C^) were detected in NL of one patient with KM-LUAD, signifying a potential field cancerization effect (**Extended Data Fig. 6c, d**). Concordantly, KRAS signature was significantly increased in *KRAS*^G12D^ KACs relative to *KRAS*^WT^ counterparts (*P* = 3.9 × 10^-3^; **Extended Data Fig. 6e**). KRAS signature was also elevated in *KRAS*^WT^ KACs relative to other AICs (*P* < 2.2 × 10^-16^) and in other AICs relative to AT2 cells (*P* < 2.2 × 10^-16^; **Extended Data Fig. 6e**), pointing towards increased KRAS signalling along the AT2-AIC-KAC spectrum. KACs from NLs or tumours of KM-but not KW-LUAD cases were consistently and significantly less differentiated than other AICs (all *P <* 2.2 × 10^-16^, **Extended Data Fig. 6f, g**). Our findings characterize KACs as an intermediate alveolar cell subset that is highly relevant to the pathogenesis of human LUAD, especially KM-LUAD.

### A KAC state is linked to mouse KM-LUAD

We next performed scRNA-seq analysis of lung epithelial cells from mice with knockout of the lung lineage-specific G-protein coupled receptor (*Gprc5a^-/-^*)^21,22^ and that form KM-LUADs following tobacco-carcinogen exposure. We analysed lungs from *Gprc5a*^-/-^ mice treated with nicotine-derived nitrosamine ketone (NNK) or control saline at the end of exposure (EOE) and at 7 months post-exposure, the time point of KM-LUAD onset (n = 4 mice per group and time point; **Fig. 3a, Supplementary Fig. 4**). Clustering analysis of 9,272 high-quality epithelial cells revealed distinct lineages including KACs that clustered in between AT1 and AT2 cell subsets and close to tumour cells (**Extended Data Fig. 7a**). Like their human counterparts, malignant cells displayed low expression of lineage-specific genes (**Extended Data Fig. 7b** and **Supplementary Table 10**). Consistently, cells from the malignant cluster had high CNV scores, expressed *Kras*^G12D^ mutations, and showed increased expression of markers associated with loss of alveolar differentiation (*Kng2* and *Meg3*) and immunosuppression (*Cd24a*^23^) (**Extended Data Fig. 7c, d**). Malignant cells were present only at 7 months post-NNK and were absent at EOE to carcinogen and in saline-treated animals (**Fig. 3b** and **Extended Data Fig. 7e, f**). KAC fractions were markedly increased at EOE relative to control saline-treated littermates (*P* = 0.03), and they were, for the most part, maintained at 7 months post-NNK (**Fig. 3b** and **Extended Data Fig. 7f, g**). Immunofluorescence (IF) analysis showed that Krt8^+^ AT2-derived cells were present in NNK-exposed NL and were nearly absent in lungs of saline-exposed mice (**Fig. 3c**). LUADs also displayed high expression of Krt8 (**Fig. 3c**). KACs displayed markedly increased prevalence of *Kras*^G12D^ mutations, more so than CNV burden, and increased expression of genes (e.g., *Gnk2*) associated with loss of alveolar differentiation^24^, albeit to lesser extents compared to malignant cells (**Fig. 3d, Extended Data Fig. 7h** and **Supplementary Table 11**). Of note, AT2 fractions were reduced with time (**Extended Data Fig. 7f, g**). ST analysis at 7 months post-NNK showed that tumour regions had significantly increased expression of *Krt8* and *Plaur* as well as spatially overlapping KAC and KRAS signatures (**Fig. 3e** and **Extended Data Fig. 8a, c, e**). In line with our human data, *Krt8* high KACs with elevated expression of KAC and KRAS signatures were enriched in “reactive”, non-neoplastic regions surrounding tumours and were themselves intermediary in the transition from normal to tumour cells (**Fig. 3e** and **Extended Data Fig. 8**).

**Fig. 3.**
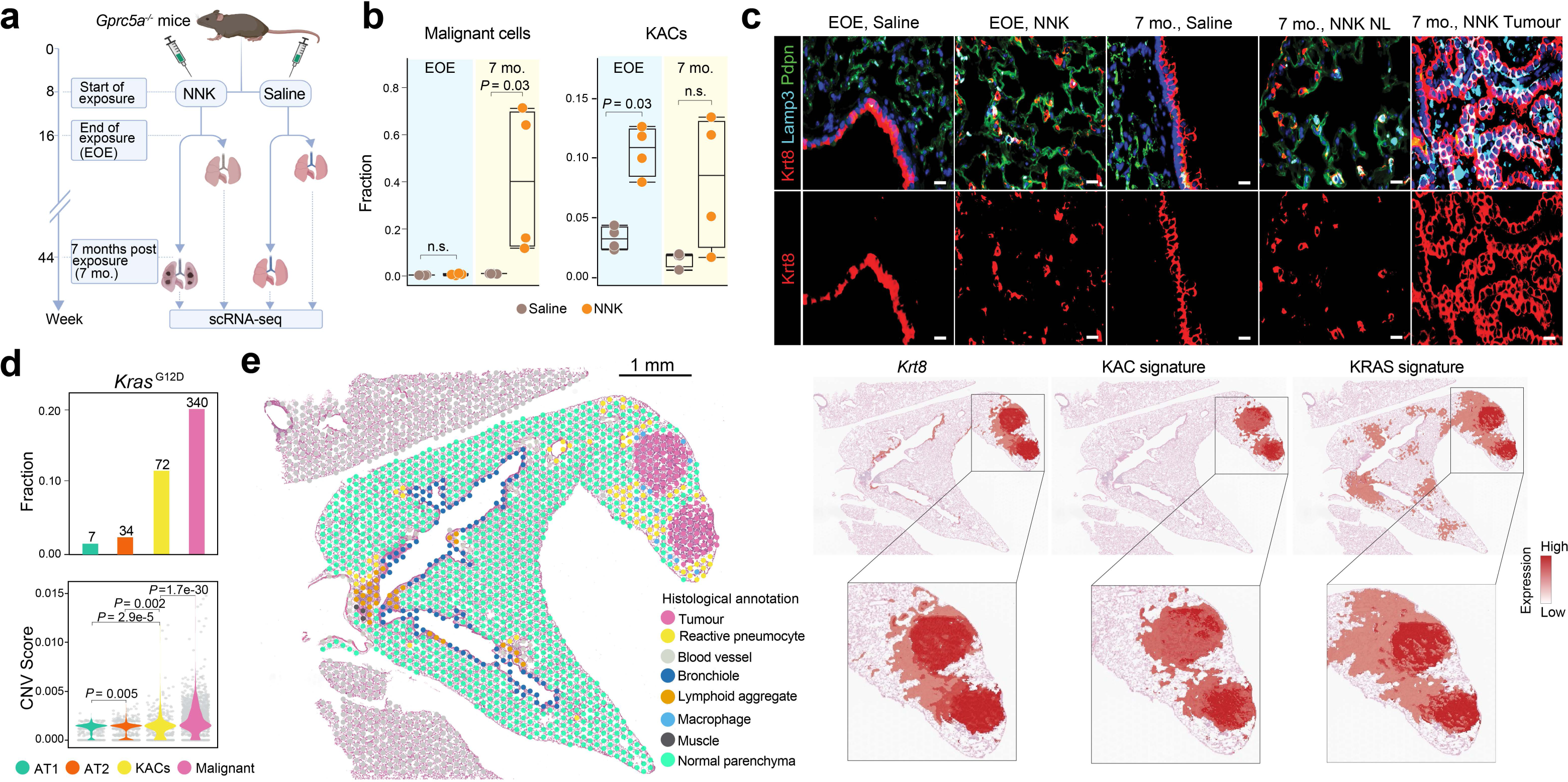
KACs evolve early and prior to tumour onset during tobacco-associated KM-LUAD pathogenesis. **a,** Schematic view of *in vivo* experimental design. mo.: months. Created with BioRender.com. **b,** Fraction of malignant cells (left) and KACs (right) across treatment groups and time points. Box-and-whisker definitions are similar to **Fig. 1g** with *n* = 4 biologically independent samples/condition. *P* - values were calculated using two-sided Mann-Whitney U test. **c,** IF analysis of KRT8, LAMP3 and PDPN in mouse lung tissues. Scale bar: 10 μm. Results are representative of two independent biological replicates per treatment and timepoint. Staining was repeated three times with similar results. **d,** Distribution of CNV scores among alveolar and malignant cells (top) and fraction of *Kras*^G12D^ mutant cells in KACs, malignant, AT1 and AT2 subsets (bottom). *n* on top of each bar denotes the numbers of *Kras*^G12D^ mutant cells in each cell group. *n* cells in lower panel: AT1 = 496; AT2 = 1,320; KACs = 512; Malignant = 1,503. *P* - values were calculated using two-sided Mann-Whitney U test with a Benjamini–Hochberg correction. **e,** ST analysis of lung tissue at 7 mo post-exposure to NNK and showing histological annotation of H&E-stained Visium slide (left) and spatial heatmaps showing scaled expression of *KRT8* as well as KAC and KRAS signatures. ST analysis was done on three different tumour-bearing mouse lung tissues from two mice at 7 months following NNK.

Mouse (**Extended Data Fig. 9a**) and human (**Extended Data Fig. 9b)** KACs displayed commonly increased activation of pathways including NF-κB, hypoxia, and p53 signalling among others. A p53 signature we derived was significantly elevated in KACs at EOE, and more so at 7 months post-exposure to NNK, compared to both AT2 as well as tumour cells (**Extended Data Fig. 9c, left)**. Similar patterns were noted for expression of p53 pathway-related genes and senescence markers including *Cdkn1a, Cdkn2b*, *Bax*, as well as *Trp53* itself (**Extended Data Fig. 9c, right)**. Of note, activation of p53 was reported in *Krt8^+^* transitional cells identified by Strunz and colleagues ^25^ during bleomycin-induced alveolar regeneration, and which themselves showed overlapping genes with KACs from our study (32%; **Extended Data Fig. 9d**). A mouse KAC signature we derived and that was significantly enriched in mouse KACs and malignant cells (*P* < 2.2 x 10^-16^, **Extended Data Fig. 9e**) and in human LUADs (*P =* 1.2 x 10^-8^, **Extended Data Fig. 9f, left**) was also significantly elevated in premalignant AAHs (*P =* 4.3 x 10^-4^) and further in invasive LUADs (*P =* 1.5 x 10^-3^) relative to matched NL tissues (**Extended Data Fig. 9f, right**). Like alveolar intermediates in acute lung injury^25,26^ and KACs in human LUADs (**Fig. 2**), mouse KACs were likely AT2-derived, acted as intermediate states in AT2 to AT1 differentiation, and also were inferred to transition to malignant cells (**Fig. 4a, top row, Supplementary Fig. 5,** and **Supplementary Table 12**). KACs assumed an intermediate differentiation state, more closely resembling malignant cells than other alveolar subsets (**Fig. 4a, middle**). KAC signature was elevated in cancer stem cell (CSC)/stem cell-like progenitor cells which we had cultured from MDA-F471 LUAD cell line (derived from an NNK-exposed *Gprc5a*^-/-^ mouse^27^) relative to parental 2D cells (**Extended Data Fig. 10a**). KACs at EOE were somewhat less differentiated relative to those at 7 months post-exposure (**Fig. 4a, bottom right**). Notably, the fraction of KACs with *Kras*^G12D^ mutation was low at EOE (∼0.02) and was increased at 7 months post-NNK (∼0.19) (**Extended Data Fig. 10b**). *Kras*^G12D^ KACs from the late timepoint were significantly less differentiated (*P* = 7.8 x 10^-6^; **Extended Data Fig. 10c**) and showed higher expression of KAC signature genes such as *Cldn4*, *Krt8*, *Cavin3*, and *Cdkn2a* relative to *Kras*^wt^ KACs (**Extended Data Fig. 10d)**. Interestingly, *Kras*^wt^ KACs were more similar to *Krt8^+^* intermediate cells from the Strunz et. al ^25^ than *Kras*^G12D^ KACs (20% overlap versus 10% respectively; **Extended Data Fig. 10e, f**).

**Fig. 4.**
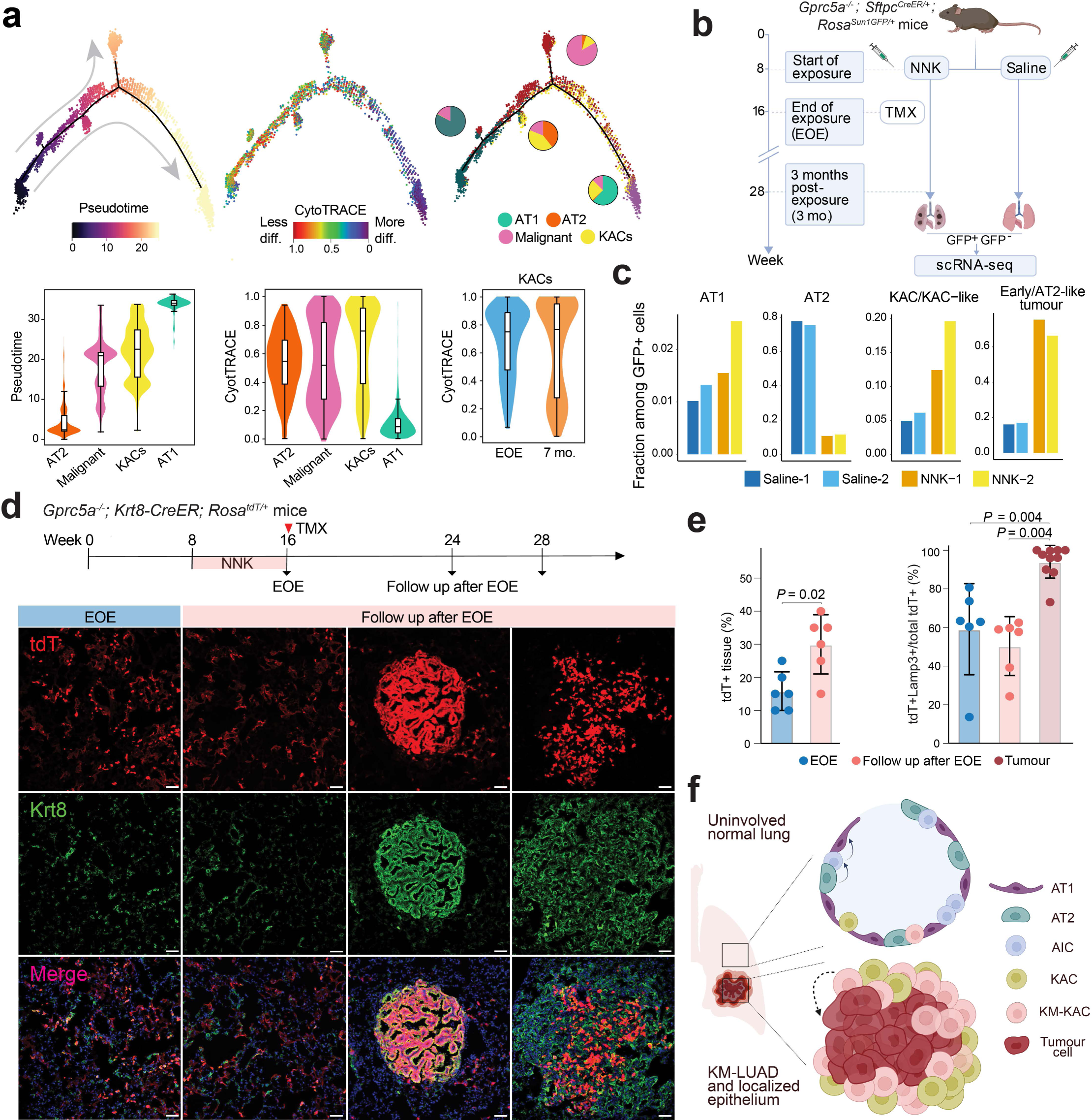
KACs are implicated in transition of AT2 to *Kras*-mutant tumour cells. **a,** Trajectories of alveolar and malignant cells coloured by inferred pseudotime, cell differentiation status, and cell type (top left to right). Distribution of inferred pseudotime (bottom left) and CytoTRACE (bottom middle) scores across the indicated cell subsets. Bottom right panel shows CytoTRACE score distribution in KACs at the two time points. Box-and-whisker definitions are similar to **Fig. 1g** with *n* cells (left to right): 1,791; 1,693; 636; 580; 1,791; 1,693; 636; 580; 301; 335. **b,** Schematic overview showing analysis of *Gprc5a^-/-^* mice with reporter labeled-AT2 cells (*Gprc5a^-/-^;* Sftpc*^CreER/+^; Rosa^Sun1GFP/+^*). TMX: tamoxifen. **c,** Fractions of AT1, AT2, KAC/KAC-like, and early/AT2-like tumour cells within GFP^+^ cells from lungs of 2 NNK and 2 saline-exposed mice analysed at 3 months post-exposure. **d,** IF analysis of tdT and Krt8 expression at EOE to NNK (first column; EOE) and at 8-12 weeks following NNK (follow up after EOE) in normal-appearing regions (second column) and tumours (last two columns) of *Gprc5a*^-/-^; Krt8-CreER; *Rosa^tdT^*^/+^ mice. Tamoxifen (1 mg/dose) was delivered right after EOE to NNK for six continuous days. Results are representative of 3 biological replicates per condition. Staining was performed two times with similar results. Magnification: 20X. Scale bar: 10 μm. **e**, Percentages of lung tissue areas containing tdT^+^ cells (left). Percentage of tdT^+^/Lamp3^+^ cells among total tdT^+^ cells in normal-appearing regions at different timepoints. Error bars show mean ± SD of *n* biologically independent samples (left to right): 6; 6; 6; 6; 10. *P* - values were calculated using Mann-Whitney U test. **f,** Proposed model for alveolar plasticity whereby a subset of AICs in the intermediate AT2-to-AT1 differentiation state are KACs and, later, acquire *KRAS*^G12D^ mutations and are implicated in KM-LUAD development from a particular region in the lung. Panels **b** and **f** were created with BioRender.com.

We performed integrated scRNA-seq analysis of cells from our mouse cohort with those in mice driven by *Kras*^G12D^ from the two separate studies by Marjanovic et al ^28^ and Dost et al ^29^. Cluster C5 comprised cells from all three studies with distinctly high expression of KAC markers and KAC signature itself (**Extended Data Fig. 10g-i**). The overwhelming fraction of C5 cells were from our study, and yet, C5 cells from *Kras*^G12D^*-*driven mice still expressed higher mouse KAC signature compared to normal AT2 cells from all studies (**Extended Data Fig. 10j**). The mouse KAC signature was markedly and significantly increased in human AT2 cells with induced expression of *KRAS*^G12D^ relative to those with wild type *KRAS* from the Dost et al study (*P* < 2.2 x 10^-16^, **Extended Data Fig. 10k**). In agreement with these findings, the mouse KAC signature, like its human counterpart (**Extended Data Fig. 4h**), was significantly enriched in KACs and malignant cells from KM-LUADs relative to EM-LUADs (*P =* 0.04 and *P* < 2.2 x 10^-16^, respectively; **Extended Data Fig. 10l**).

We further investigated the biology of KACs using *Gprc5a^-/-^*mice with reporter labeled-AT2 cells (*Gprc5a^-/-^; S*ftpc*^CreER/+^; Rosa^Sun1GFP/+^*; **Fig. 4b**). GFP^+^ organoids derived from NNK-but not saline-exposed reporter mice at EOE were enriched with KACs (**Extended Data Fig. 11a, Supplementary Fig. 6**). GFP^+^ cells (n = 3,089) almost exclusively comprised AT2, early/AT2-like tumour cells, KAC/KAC-like cells, and few AT1 cells, all of which were nearly absent in the GFP^-^ fraction (**Extended Data Fig. 11b, c and Supplementary Fig. 7**). There were markedly increased fractions of GFP^+^ AT1, KACs, and, expectedly, tumour cells from NNK-versus saline-treated mice (**Fig. 4c**). GFP expression was almost exclusive to alveolar regions and tumours, the latter which were almost entirely GFP^+^ as well as Krt8^+^ and KAC marker^+^ (Cldn4^+^, Cavin3^+^) (**Supplementary Fig. 8a-c**). Normal lung regions included AT2-derived KACs (GFP^+^/Krt8^+^ and Cldn4^+^ or Cavin3^+^) (**Supplementary Fig. 8a-c**). GFP^+^/Lamp3^+^/Krt8^-/low^ AT2 cells were also evident including in normal (non-tumoral) lung regions from NNK-exposed reporter mice (**Supplementary Fig. 8d**). GFP^+^ KACs from this time point that coincides with formation of preneoplasias ^21^ harboured driver *Kras*^G12D^ mutations at similar fractions when compared with early/AT2-like tumour cells (**Extended Data Fig. 11d-f)**. As seen in *Gprc5a^-/-^*mice (**Fig. 4a**), KACs were closely associated with tumour cells in pseudotime (**Extended Data Fig. 11g, h**).

GFP^+^ organoids from reporter mice at 3 months post-NNK showed significantly and markedly enhanced growth compared to those from saline-exposed animals and were almost exclusively comprised of cells with KAC markers (Krt8^+^, Cldn4^+^; **Extended Data Fig. 12a, e**). Given that KACs, like early tumour cells, acquired *Kras* mutations, we examined effects of targeted KRAS^G12D^ inhibition on these organoids. We first tested effects of KRAS^G12D^ inhibitor MRTX1133^30^ *in vitro* and found that it inhibited the growth of mouse MDA-F471 as well as LKR13 (derived from *Kras^LSL-G12D^* mice; ^31^) cells in a dose-dependent manner (**Extended Data Fig. 12b)**. This was accompanied by suppression of phosphorylated levels of ERK1/2 and S6 kinase in both cell lines (**Extended Data Fig. 12c and Supplementary Fig. 9**). Notably, MRTX1133-treated KAC marker positive organoids showed significantly reduced sizes as well as Krt8 and Cldn4 expression intensities relative to DMSO-treated counterparts (*P* < 1.5 x 10^-10^; **Extended Data Fig. 12d, e**).

To further confirm that KACs indeed give rise to tumour cells, we labelled Krt8^+^ cells in *Gprc5a^-/-^;* Krt8-CreER*; Rosa^tdT^*^/+^ mice. Krt8-CreER; *Rosa^tdT^*^/+^ mice were first used to confirm increased tdT^+^ labelling (i.e., higher number of KACs) in the lung parenchyma at EOE to NNK compared to control saline-treated mice (**Fig. 4d** and **Extended Data Fig. 13a**). We then analysed lungs of NNK-exposed *Gprc5a^-/-^; Krt*8-CreER; *Rosa^tdT^*^/+^ mice that were injected with tamoxifen right after completing NNK (**Fig. 4d**). Of note, most tumours showed tdT^+^/Krt8^+^ cells at varying levels, with some tumours showing strong extent of tdT labelling and suggesting oncogenesis of Krt8^+^ cells (**Fig. 4d** and **Extended Data Fig. 13b, c**). Most tdT^+^ tumour cells were AT2 derived (Lamp3^+^) (**Fig. 4e** and **Extended Data Fig. 13b**). Fraction of tdT^+^/Lamp3^+^ cells out of total tdT^+^ cells was similar between EOE and follow up after EOE to NNK (**Fig. 4e**). Normal-appearing regions also showed tdT^+^ AT1 cells (Nkx2-1^+^/Lamp3^-^) suggesting possible turnover of AT2 cells and KACs to AT1 cells (**Extended Data Fig. 13a**). Taken together, our *in vivo* analyses identify KACs as an intermediate cell state in early development of KM-LUAD and following tobacco carcinogen exposure.

## Discussion

Our multi-modal analysis of epithelial cells from early-stage LUADs and the peripheral lung uncovered diverse malignant states, patterns of ITH, and cell plasticity programs that are linked to KM-LUAD pathogenesis. Of these, we identified alveolar intermediary cells (KACs) that arise upon activation of alveolar differentiation programs and that could act as progenitors for KM-LUAD (**Fig. 4f**). KACs were evident in normal-appearing areas in the vicinity of lesions in both mouse and patient samples, suggesting that the early appearance of these cells (e.g., following tobacco exposure) may represent a *field of injury*^11^. A pervasive field of injury is relevant to development of human lung cancer and to the complex spectrum of mutations present in normal-appearing lung tissue ^32,33^. We propose that KACs represent injured/mutated cells in the normal-appearing lung that have increased likelihood for transformation to lung tumour cells (**Fig. 4f**).

Our analysis uncovered strong links and intimately shared properties between KACs and *KRAS*- mutant lung tumour cells including *KRAS* mutations, reduced differentiation, and pathways.

Notably, we show that growth of KAC-rich and AT2 reporter-labelled organoids derived from lungs with early lesions was highly sensitive to KRAS^G12D^ inhibition^34^. While our *in vivo* findings are consistent with previous independent reports showing that AT2 cells are the preferential cells-of-origin in *Kras*-driven LUADs in animals^35–37^, they enable a deeper scrutiny of the unique attributes and states of alveolar intermediary cells in the trajectory towards KM-LUADs.

Following acute lung injury, AT2 cells were shown to differentiate into AICs that are characterized by high expression of *Krt8* and that are crucial for AT1 regeneration ^25,26,38^. We did find evidence of KAC-like cells with notable expression of the KAC signature in *Kras*^G12D^-driven mice, albeit at lesser frequency compared to our tobacco-mediated carcinogenesis model. Thus, it is plausible that KACs can arise due to an injury stimulus (here tobacco exposure) or mutant *Kras* expression – or due to both. Our work introduces open questions that would be important to pursue in future studies. It is not clear whether KACs are a dominant or obligatory path in AT2-to-tumour transformation. Also, we do not know the effects of expressing mutant oncogenes, *Kras* or others, or tumour suppressors on the likelihood of KACs to divert away from mediating AT1 regeneration and, instead, transition to tumour cells. Recent studies suggest that p53 could curtail the oncogenesis of alveolar intermediate cells ^39^.

Our study with a marriage between in-depth interrogation of early-stage human LUADs and *Kras*-mutant lung carcinogenesis models provides an atlas with an expansive number of epithelial cells. This atlas of epithelial and malignant cell states in human and mouse lungs underscores new cell-specific subsets that underlie inception of LUADs. Our discoveries inspire the derivation of targets (e.g., KAC signals such as early KRAS programs) to prevent initiation and development of LUAD.

## Methods

### Multi-regional sampling of human surgically resected LUADs and normal lung tissues

Study subjects were evaluated at MD Anderson Cancer Center and underwent standard-of-care surgical resection of early-stage LUAD (I–IIIA). Samples from all patients were obtained from banked or residual tissues under informed consents and approved by MD Anderson institutional review board (IRB) protocols. Residual surgical specimens were then used for derivation of multi-regional samples for single-cell analysis (**Supplementary Table 1**). Immediately following surgery, resected tissues were processed by an experienced pathologist assistant. One side of the specimen was documented and measured, followed by tumour margin identification. Based on placement of the tumour within the specimen, incisions were made at defined collection sites in one direction along the length of the specimen and spanning the entire lobe: tumour-adjacent and tumour-distant normal parenchyma at 0.5 cm from the tumour edge and from the periphery of the overall specimen/lobe, respectively. An additional tumour-intermediate normal tissue was selected for P2–P16 and ranged between 3 and 5 cm from the edge of the tumour. Sample collection was initiated with normal lung tissues that are farthest from the tumour moving inward toward the tumour to minimize cross-contamination during collection.

### Single-cell isolation from tissue samples

Fresh tissues from human donors as well as mouse lungs were freshly collected in RPMI medium supplemented with 2% foetal bovine serum (FBS) and maintained on ice for immediate processing. Tissues were placed in a cell culture dish containing Hank’s balanced salt solution (HBSS) on ice and extra-pulmonary airways and connective tissue were removed with scissors. Samples were transferred to a new dish on ice and minced into ∼1mm^3^ pieces followed by enzymatic digestion using a cocktail for human tissues composed of Collagenase A (10103578001, Sigma Aldrich), Collagenase IV (NC9836075, Thermo Fisher Scientific), DNase I, (11284932001, Sigma Aldrich), Dispase II (4942078001, Sigma Aldrich), Elastase (NC9301601, Thermo Fisher Scientific) and Pronase (10165921001, Sigma Aldrich) as previously described ^40^, or a cocktail for mouse lung digestion composed of Collagenase Type I (CLS-1 LS004197, Worthington), Elastase (ESL LS002294, Worthington), and DNase I (D LS002007, Worthington). Samples were transferred to 5 ml LoBind Eppendorf tubes and incubated in a 37°C oven for 20 minutes with gentle rotation. Samples were then filtered through 70 μm strainers (Miltenyi biotech, 130-098-462) and washed with ice-cold HBSS. Filtrates were then centrifuged and resuspended in ice-cold ACK lysis buffer (A1049201, Thermo Fisher Scientific) for red blood cell (RBC) lysis. Following RBC lysis, samples were centrifuged and resuspended in ice-cold FBS, filtered (using 40 μm FlowMi tip filters; H13680-0040, Millipore), and an aliquot was taken to count cells and check for viability by Trypan blue (T8154, Sigma Aldrich) exclusion analysis.

### Sorting and enrichment of viable lung epithelial singlets

Single cells from patient 1 (P1) were stained with SYTOX Blue viability dye (S34857, Life Technologies), and processed on a fluorescence-activated cell sorting (FACS) Aria I instrument. Cells from patients 2 through 16 (P2-P16) were stained with anti-EPCAM-PE (347198, BD Biosciences;1:50 dilution in ice-cold phosphate-buffered saline, PBS, containing 2% FBS) for 30 minutes with gentle rotation at 4°C. Mouse lung single cells were similarly stained but with a cocktail of antibodies (1:250 each) against CD45-PE/Cy7 (103114, BioLegend), Icam2-A647 (A15452, Life Technologies), Epcam-BV421 (118225, BioLegend), and Ecad-A488 (53-3249-80, eBioscience). Stained cells were then washed, filtered using 40 μm filters, stained with SYTOX Blue (human) or SYTOX Green (mouse) and processed on a FACS Aria I instrument (gating strategies for epithelial cell sorting are shown in **Supplementary Fig. 1 and 4 for human and mouse cells, respectively**). Doublets and dead cells were eliminated, and viable (SYTOX-negative) epithelial singlets were collected in PBS containing 2% FBS. Cells were washed again to eliminate ambient RNA, and a sample was taken for counting by Trypan Blue exclusion before loading on 10X Genomics Chromium microfluidic chips.

### Preparation of single-cell 5’ gene expression libraries

Up to 10,000 cells per sample were partitioned into nanolitre-scale Gel beads-in-emulsion (GEMs) using Chromium Next GEM Single Cell 5’ Gel Bead kit v1.1 (1000169, 10X Genomics, Pleasanton, CA) and by loading onto Chromium Next GEM Chips G (1000127, 10X Genomics, Pleasanton, CA). GEMs were then recovered to construct single-cell gene expression libraries using the Chromium Next GEM Single Cell 5’ Library kit (1000166, 10X Genomics) according to the manufacturer’s protocol. Briefly, recovered barcoded GEMs were broken and pooled, followed by magnetic bead clean-up (Dynabeads MyOne Silane, 37002D, Thermo Fisher Scientific). 10X-barcoded full-length cDNA was then amplified by PCR and analysed using Bioanalyzer High Sensitivity DNA kit (5067-4626, Agilent). Up to 50 ng of cDNA was carried over to construct gene expression libraries and was enzymatically fragmented and size-selected to optimize the cDNA amplicon size prior to 5’ gene expression library construction. Further, samples were subject to end-repair, A-tailing, adaptor ligation, and sample index PCR using Single Index kit T Set A (2000240, 10X Genomics) to generate Illumina-ready barcoded gene expression libraries. Library quality and yield was measured using Bioanalyzer High Sensitivity DNA (5067-4626, Agilent) and Qubit dsDNA High Sensitivity Assay (Q32854, Thermo Fisher Scientific) kits. Indexed libraries were normalized by adjusting for the ratio of the targeted cells per library as well as individual library concentration and then pooled to a final concentration of 10 nM. Library pools were then denatured and diluted as recommended for sequencing on the Illumina NovaSeq 6000 platform.

### Single-cell RNA-seq data processing and quality control

Raw scRNA-seq data were pre-processed (demultiplex cellular barcodes, read alignment, and generation of gene count matrix) using Cell Ranger Single Cell Software Suite (version 3.0.1) provided by 10X Genomics. For read alignment of human and mouse scRNA-seq data, human reference GRCh38 (hg38) and mouse reference GRCm38 (mm10) genomes were used, respectively. Detailed quality control (QC) metrics were generated and evaluated, and cells were carefully and rigorously filtered to obtain high-quality data for downstream analyses^15^. Briefly, for basic quality filtering, cells with low complexity libraries (in which detected transcripts were aligned to < 200 genes such as cell debris, empty drops, and low-quality cells) were filtered out and excluded from subsequent analyses. Likely dying or apoptotic cells in which > 15% of transcripts derived from the mitochondrial genome were also excluded. For scRNA-seq analysis of *Gprc5a^-/-^*; Sftpc*^CreER/+^*; *Rosa^Sun1GFP/+^* mice, cells with ≤ 500 detected genes or with mitochondrial gene fraction that is ≥ 15% were filtered out using Seurat^41^.

### Doublet detection and removal, and batch effect evaluation and correction

Likely doublets or multiplets were identified and carefully removed through a multi-step approach as described in our recent studies^15,42^. Briefly, doublets or multiplets were identified based on library complexity, whereby cells with high-complexity libraries in which detected transcripts are aligned to > 6,500 genes were removed, and also, based on cluster distribution and marker gene expression, whereby doublets or multiplets forming distinct clusters with hybrid expression features and/or exhibiting an aberrantly high gene count were also removed. Expression levels and proportions of canonical lineage-related marker genes in each identified cluster were carefully reviewed. Clusters co-expressing discrepant lineage markers were identified and removed. Doublets or multiplets were also identified using the doublet detection algorithm DoubletFinder^43^. The proportion of expected doublets was estimated based on cell counts obtained prior to scRNA-seq library construction. Data normalization was then performed using Seurat^41^ on the filtered gene-cell matrix. Statistical assessment of possible batch effects was performed on non-malignant epithelial cells using the R package ROGUE^36^, an entropy-based statistic, as described in our previous studies^15,42^ and Harmony ^44^ was run with default parameters to remove batch effects present in the PCA space.

### Unsupervised clustering and subclustering analysis

The function *FindVariableFeatures* of Seurat^41^ was applied to identify highly variable genes (HVGs) for unsupervised cell clustering. PCA was performed on the top 2,000 HVGs. The elbow plot was generated with the *ElbowPlot* function of Seurat and, based on which, the number of significant principal components (PCs) was determined. The *FindNeighbors* function of Seurat was used to construct the Shared Nearest Neighbour (SNN) graph, based on unsupervised clustering performed with Seurat function *FindClusters*. Multiple rounds of clustering and subclustering analysis were performed to identify major epithelial cell types and distinct cell transcriptional states. Dimensionality reduction and 2-D visualization of cell clusters was performed using UMAP^45^ with Seurat function *RunUMAP*. The number of PCs used to calculate the embedding was the same as that used for clustering. For analysis of human epithelial cells, ROGUE was employed to quantify cellular transcriptional heterogeneity of each cluster. Subclustering analysis was then performed for low-purity clusters identified by ROGUE. Hierarchical clustering of major epithelial subsets was performed on the Harmony batch corrected PCA dimension reduction space. For malignant cells, except for global UMAP visualization, downstream analyses including identification of large-scale copy number variations (CNVs), inference of cancer cell differentiation states, quantification of meta-program expressions, trajectory analysis, and mutation analysis were performed without Harmony batch correction. The hierarchical tree of human epithelial cell lineages was computed based on Euclidean distance with Ward linkage method and the dendrogram was generated using the R function *plot.hc*. For scRNA-seq analysis of *Gprc5a*^−/−^ mice, the top-ranked 10 PCs were selected using the *elbowplot* function. SNN graph construction was performed with resolution parameter = 0.4 and UMAP visualization was performed with default parameters. For scRNA-seq analysis of *Gprc5a^-/-^*; *S*ftpc*^CreER/+^*; *Rosa^Sun1GFP/+^* mice, the top-ranked 20 Harmony-corrected PCs were used for SNN graph construction and unsupervised clustering was performed with resolution parameter = 0.4. UMAP visualization was performed with *RunUMAP* function with min.dist = 0.1. Differentially expressed genes (DEGs) of clusters were identified using *FindAllMarkers* function with FDR adjusted *P* - value < 0.05 and log2 fold change > 1.2.

### Identification of malignant cells and mapping *KRAS* codon 12 mutations

Malignant cells were distinguished from non-malignant subsets based on information integrated from multiple sources as described in our recent studies^15,42^. The following strategies were used to identify malignant cells: ***cluster distribution***: due to the high degree of inter-patient tumour heterogeneity, malignant cells often exhibit distinct cluster distribution from that of normal epithelial cells. While non-malignant cells derived from different patients often are clustered together by cell type, malignant cells from different patients likely form separate clusters. ***CNVs***: We applied inferCNV^16^ (version 1.3.2) to infer large-scale CNVs in each individual cell with T cells as the reference control. To quantify copy number variations at the cell level, CNV scores were aggregated using a strategy like that described in a previous study^16^. Briefly, arm-level CNV scores were computed based on the mean of the squares of CNV values across each chromosomal arm. Arm-level CNV scores were further aggregated across all chromosomal arms by calculating the arithmetic mean value of the arm-level scores using R function *mean*. ***Marker gene expression***: expression of lung epithelial lineage-specific genes and LUAD-related oncogenes was determined in epithelial cell clusters. ***Cell-level expression of KRAS*^G12D^ *mutations***: as we previously described ^15^, BAM files were queried for *KRAS*^G12D^ mutant alleles which were then mapped to specific cells. *KRAS*^G12D^ mutations, along with cluster distribution, marker gene expression, and inferred CNVs as described above, were used to distinguish malignant cells from non-malignant cells. Following clustering of malignant cells from all patients, an absence of malignant cells that were identified from P12 or P16 was noted. This can be possibly attributed to the low number of epithelial cells captured in tumour samples from these patients (**Supplementary Table 2**).

### Mapping KRAS codon 12 mutations

To map somatic *KRAS* mutations at single cell resolution, alignment records were extracted from the corresponding BAM files using mutation location information. Unique mapping alignments (MAPQ=255) labelled as either PCR duplication or secondary mapping were filtered out. The resulting somatic variant carrying reads were evaluated using Integrative Genomics Viewer (IGV)^46^ and the “CB” tags were used to identify cell identities of mutation-carrying reads. To estimate the variant allele fraction (VAF) of *KRAS*^G12D^ mutation and cell fraction of *KRAS*^G12D^-carrying cells within malignant and non-malignant epithelial cell subpopulations (e.g., malignant cells from all LUADs, malignant cells from KM-LUADs, KACs from KM-LUADs), reads were first extracted based on their unique cell barcodes and bam files were generated for each subpopulation using samtools (v1.15). Mutations were then visualized using IGV, and VAFs were calculated by dividing the number of *KRAS*^G12D*-*^carrying reads by the total number of uniquely aligned reads for each subpopulation. A similar approach was used to visualize *KRAS*^G12C^*-*carrying reads and to calculate the VAF of *KRAS*^G12C^ in KACs of normal tissues from KM cases. To calculate the mutation-carrying cell fraction, extracted reads were mapped to the *KRAS*^G12D^ locus from bam files using *AlignmentFile* and *fetch* functions in *pysam* package. Extracted reads were further filtered using the ‘Duplicate’ and ‘Quality’ tags to remove PCR duplicates and low-quality mappings. The number of reads with/without *KRAS*^G12D^ mutation in each cell was summarized using the ‘CB’ tag in read barcodes. Mutation-carrying cell fractions were then calculated as the ratio of the number of cells with at least one *KRAS*^G12D^ read over the number of cells with at least one high-quality read mapped to the locus.

### PCA analysis of malignant cells and quantification of transcriptome similarity

Raw unique molecular identifier (UMI) counts of identified malignant cells were log normalized and used for PCA analysis using Seurat (*RunPCA* function). PCA dimension reduction data were extracted using *Embeddings* function. The top three most highly ranked PCs were exported for visualization using JMP v15. 3D scatterplots of PCA data were generated using the scatterplot 3D tool in JMP v15. Bhattacharyya distances were calculated using the *bhattacharyya.dist* function from R package fpc (v2.2-9). Top 25 highly ranked PCs were used for both patient level and cell level distance calculations. For Bhattacharyya distance quantification at the cell level, 100 cells were randomly for each patient group defined by driver mutations (e.g., KM-LUADs). The random sampling process was repeated 100 times and pairwise Bhattacharyya distances were then calculated between patient groups. Differences in Bhattacharyya distances between patient groups were tested using Wilcoxon Rank-Sum tests and boxplots were generated using the *geom_boxplot* function from R package ggplot2 (v3.2.0).

### Determination of non-malignant cell types and states

Non-malignant cell types and states were determined based on unsupervised clustering analysis following batch effect correction using Harmony^44^. Two rounds of clustering analysis were performed on non-malignant cells to identify major cell types and cell transcriptional states within major cell types. Clustering and UMAP visualization of human normal epithelial cells (**Extended Data Fig. 1a**) was performed using Seurat with default parameters. Specifically, the parameters k.param = 20 and resolution = 0.4 were used for SNN graph construction and cluster identification, respectively. UMAP visualization was performed with default parameters (min.dist = 0.3). For clustering analysis of airway and alveolar epithelial cells, *RunPCA* function was used to determine the most contributing top PCs for each subpopulation and similar clustering parameters (k.param = 20 and resolution = 0.4) were used for SNN graph construction and cluster identification. UMAP plots were generated with min.dist = 0.3 using the *RunUMAP* function in Seurat. Density plots of alveolar intermediate cells were generated using the *stat_densit_2d* function in R package ggplot2 (version: 3.3.5) with the first two UMAP dimension reduction data as the input. DEGs for each cluster were identified using the *FindMarkers* function in Seurat with an FDR adjusted *P* < 0.05 and a fold change cut-off > 1.2. Canonical epithelial marker genes from previously published studies by our group and others ^15,47,48^ were used to annotate normal epithelial cell types and states. Bubble plots were generated for select DEGs and canonical markers to define AT1 (*AGER1*/*ETV5/PDPN*+), AT2 (*SFTPB*/*SFTPC*/*ETV5*+), SDP (*SCGB1A1/SFTPC* dual positive cells), AIC (*AGER1*/*ETV5/PDPN+ and SFTPB*/*SFTPC+*), club and secretory (*SCGB1A1*/*SCGB3A1/CYP2F1+*), basal (*KRT5*/*TP63*+), ciliated (*CAPS*/*PIFO*/*FOXJ1*+), ionocyte (*ASCL3*/*FOXI1*+), neuroendocrine (*CALCA*/*ASCL1*+) and tuft (*ASCL2*/*MGST2*/*PTGS1*+) cells. KACs were identified by unsupervised clustering of AICs and defined based on previously reported marker genes^25,26,49^ including significant upregulation of the following genes relative to other alveolar cells: *KRT8*, *CLDN4*, *PLAUR* as well as *CDKN1A*, and *CDKN2A*.

### Scoring of curated gene signatures

Genes in previously defined ITH meta-programs (MPs) by Gavish *et al*. ^19^ were downloaded from the original study. Among a total of 41 consensus ITH MPs identified, MPs with unassigned functional annotations (unassigned MPs 38-41; n = 4), neural/ hematopoietic lineage specific MPs (MPs 25-29, MPs 33-37; n = 10) as well as cell type-specific MPs irrelevant to LUAD (MPs 22-24 secreted/cilia, MP 32 skin-pigmentation; n = 4) were filtered out, resulting in 23 MPs which closely correlated with hallmarks of cancer and which were used for further analysis. Signature scores were computed using the *AddModuleScore* function in *Seurat* as described previously^15,42^. KRAS signature used in this study was derived by calculating DEGs between the *KRAS*-mutant malignant cell-enriched cluster and other malignant cells (FDR adjusted *P* - value < 0.05, log fold change > 1.2, **Extended Data Fig. 2i**). Human and mouse KAC signatures as well as human “other AIC” signature were derived by calculating DEGs using *FindAllMarkers* among alveolar cells (FDR adjusted *P* - value < 0.05, log fold change > 1.2). Mouse genes in the tp53 pathway were downloaded from the Molecular Signature Database (MSigDB: https://www.gsea-msigdb.orggsea/msigdb/mouse/geneset/HALLMARK_P53_PATHWAY; MM3896). Signature scores for KACs, other AICs, KRAS and tp53 were calculated using the *AddModuleScore* function in Seurat.

### Analysis of alveolar cell differentiation states and trajectories

Analysis of differentiation trajectories of lung alveolar and malignant cells was performed using Monocle 2^50^ by inferring the pseudotemporal ordering of cells according to their transcriptome similarity. Monocle 2 analysis of malignant cells from P14 was performed with default parameters using the *detectGenes* function. Detected genes were further required to be expressed by at least 50 cells. For construction of the differentiation trajectory of lineage-labelled epithelial cells (GFP^+^), the top 150 DEGs (FDR adjusted *P* - value < 0.05, log fold change > 1.5, expressed in ≥ 50 cells) ranked by fold-change of each cell population from NNK-treated samples were used for ordering cells with setOrderingFilter function. Trajectories were generated using the reduceDimension function with method set to ‘DDRTree’. Trajectory roots were selected based on 1) inferred pseudotemporal gradient and 2) CytoTRACE score prediction and 3) careful manual review of the DEGs along the trajectory. To depict expression dynamics of ITH MPs ^19^, ITH MP scores were plotted along the pseudotime axis and smoothed lines were generated using the *smoother* tool in JMP Pro v15. Using the raw counts without normalization as input, CytoTRACE^18^ was applied with default parameters to infer cellular differentiation states to complement trajectory analysis and further understand cellular differentiation hierarchies. The *normalmixEM* function from mixtools R package was used to determine the CytoTRACE score threshold in AICs with k = 2. A final threshold of 0.58 was used to dichotomize AICs into high and low differentiation groups. Wasserstein distance metric was applied using R package transport v0.13 to quantify the variability of distribution of CytoTRACE scores. Function *wasserstein1d* was used to calculate the distance between the distribution of actual CytoTRACE scores of one patient and the distribution of simulated data with identical mean and standard deviation. The robustness of Monocle 2-based pseudotemporal ordering prediction was validated by independent pseudotime prediction tools including Palantir ^51^ , Slingshot ^52^ and Cellrank ^53^. Slingshot (v2.6.0) pseudotime prediction was performed using *slingshot* function with reduceDim parameter set to ‘PCA’ and other parameters set to defaults. Cellrank prediction was performed using the *CytoTRACEKernel* function with default parameters from Cellrank python package (v1.5.1). Palantir prediction was performed using Palantir python package (v1.0.1). A diffusion map was generated using *run_diffusion_maps* function with n_components = 5. Palantir prediction was generated using run_*palantir* function with num_waypoints = 500 and other parameters set to defaults. Inferred pseudotime by the three independent methods was then integrated with that generated by Monocle 2 for each single cell, followed by pairwise mapping and correlation analysis. Cell density plots were generated using *Contour* tool in JMP v15 with n = 10 gradient levels and contour type parameter set to ‘Nonpar Density’. To assess the pseudotime prediction consistency between Monocle 2 and the three independent methods, Spearman’s correlation coefficients were calculated and statistically tested using *cor*.*test* function in R.

### Spatial transcriptomics data generation and analysis

Spatial transcriptomic (ST) profiling of FFPE tissues of P14 LUAD sample and of three lung tissues from two *Gprc5a^-/-^* mice was performed using the Visium platform from 10X Genomics according to the manufacturer’s recommendations and as previously reported by our group^54^. FFPE tissues were collected from areas adjacent to the tissues analysed by scRNA-seq. Regions of interest per tissue/sample, each comprising a 6.5 x 6.5 mm capture area, were selected based on careful annotation of H&E-stained slides that were digitally acquired using the Aperio ScanScope Turbo slide scanner (Leica Microsystems Inc., Buffalo Grove, IL). HALO software (Indica Labs, Albuquerque, NM) was used for pathological annotation (tumour areas, blood vessels, bronchioles, lymphoid cell aggregates, macrophages, muscle tissue, normal parenchyma, and reactive pneumocytes) of H&E histology images. Spot-level histopathological annotation and visualization was generated using loupe browser (v6.3.0). Briefly, cloupe files generated from Space Ranger were loaded into the loupe browser. Visualization of annotation was then generated in svg formats using the *export plot* tool. Spatial transcriptomic RNA-seq libraries were generated according to manufacturer’s instructions each with up to ∼3,600 uniquely barcoded spots. Libraries were sequenced on the Illumina NovaSeq 6000 platform to achieve a depth of at least 50,000 mean read pairs/spot and at least 2,000 median genes/spot.

Demultiplexed raw sequencing data were aligned and gene level expression quantification was generated with analysis pipelines we previously employed^54^. Briefly, demultiplexed clean reads were aligned against the UCSC human GRCh38 (hg38) or the GRCm38 (mm10) mouse reference genomes by Spaceranger (version 1.3.0 for human ST data and version 2.0.0 for mouse ST data) and using default settings. Generated ST gene expression count matrices were then analysed using Seurat v4.1.0 to perform unsupervised clustering analysis. Using default parameters, the top-ranked 30 PCA components were employed for SNN graph construction and clustering as well as UMAP low dimension space embedding with default parameters. UMAP analysis was performed using the *RunUMAP* function. The *SpatialDimPlot* function was used to visualize unsupervised clustering. The inferCNV ^16^ R package was used for copy number analysis. Reference spots used in CNV analysis were selected based on 1) careful review of cluster marker genes using the *DotPlot* function from Seurat and 2) inspection of pathological annotation. CNV scores were calculated by computing the standard deviations of copy number variations inferred across 22 autosomes. Loupe browser (v6.3.0) was used for visualization of pathological annotation results. Expression levels of genes of interest (e.g., *KRT8*) as well as signatures of interest (e.g., KAC, KRAS) were measured and directly annotated on histology images with pixel level resolution using the TESLA (v1.2.2) machine learning framework (https://github.com/jianhuupenn/TESLA ^55^). TESLA can compute superpixel-level gene expression and detect unique structures within and surrounding tumours by integrating information from high-resolution histology images. The *annotation* and *visualize_annotation* functions were used to annotate regions with high signature signals. “*KRT8*”, “*PLAUR*”, “*CLDN4*”, “*CDKN1A”*, and “*CDKN2A*” were used for “KAC markers” signature annotation in the human ST analysis. For mouse ST data, “*Krt8*”, “*Plaur*”, “*Cldn4*” and “*Cdkn1a*”, “*Cdkn2a*” were used for “KAC signature” annotation. Gene level expression visualization of “*Krt8”* and “*Plaur”* was generated using the *scatter* function from scanpy (v1.9.1). Deconvolution analysis was conducted using CytoSPACE (https://github.com/digitalcytometry/cytospace ^56^). Annotated scRNA-seq data were first transformed into a compatible format using function *generate_cytospace_from_scRNA_seurat_object*. Visium spatial data were prepared using function *generate_cytospace_from_ST_seurat_object*. Deconvolution was performed using *CytoSpace* function (v1.0.4) with default parameters. To determine neighbouring cell composition for a specific cell population in Visium data, CytoSPACE was first applied to annotate every spot with most probable cell type. Neighbouring spots were defined as the six spots surrounding each spot and, accordingly, the neighbouring cell composition for specific cell types were computed. Trajectory construction of ST data was performed using Monocle 2^18^ with the DDRTree method using DEGs with FDR adjusted *P* - value < 0.05.

### Bulk DNA extraction and WES

Total DNA was isolated from homogenized cryosections of human lung tissues and, when available, from frozen peripheral blood mononuclear cells (PBMCs) using the Qiagen AllPrep mini kit (80204) or DNeasy Blood & Tissue kit (69504), respectively (both from Qiagen, Germantown, MD) according to the manufacturer’s recommendations. Qubit 4 Fluorometer (Thermo Fisher Scientific, Waltham, MA) was used for measurement of DNA yield. TWIST-WES was performed on the NovaSeq 6000 platform at a depth of 200X for tumour samples and 100X for NL and PBMCs to analyse recurrent driver mutations and using either PBMCs, or distant NL tissues when blood draw was not consented, as germline control. WES data were processed and mapped to human reference genome and somatic mutations were identified and annotated as previously described^57,58^ with further filtration steps. Briefly, only MuTect^59^ calls marked as “KEEP” were selected and taken into the next step. Mutations with a low variant allelic fraction (VAF < 0.02) or low alt allele read coverage (< 4) were removed. Then, common variants reported by ExAc (the Exome Aggregation Consortium, http://exac.broadinstitute.org), Phase-3 1000 Genome Project (http://phase3browser.1000genomes.org/Homo_sapiens/Info/Index), or the NHLBI GO Exome Sequencing Project (ESP6500) (http://evs.gs.washington.edu/EVS/) with minor allele frequencies greater than 0.5% were further removed. Intronic mutations, mutations at 3’ or 5’ UTR or UTR flanking regions, and silent mutations were also removed. The mutation load in each tumour was calculated as the number of nonsynonymous somatic mutations (nonsense, missense, splicing, stop gain, stop loss substitutions as well as frameshift insertions and deletions).

### Survival analysis

Analysis of overall survival (OS) in the TCGA LUAD and PROSPECT ^60^ cohorts was performed as previously described^15^. *KRAS* mutation status in TCGA LUAD samples was downloaded from cBioPortal (https://www.cbioportal.org, study ID: luad_tcga_pan_can_atlas_2018). For the TCGA dataset, clinical data were downloaded from the PanCanAtlas study^18^. Logrank test and Kaplan–Meier method were used to calculate *P* - values between groups and to generate survival curves, respectively. Statistical significance testing for all survival analyses was two-sided. To control for multiple hypothesis testing, Benjamini–Hochberg method was applied to correct *P* - values and FDR *q* - values were calculated where applicable. Results were considered statistically significant at *P* - value or FDR *q* - value of < 0.05. Multivariate survival analysis was performed using a Cox proportional hazards (PH) regression model that calculated the Hazard Ratio (HR), the 95% confidence interval (95% CI), and *P* values when using pathologic stage, age, KAC and “other AIC” signatures as covariables.

### Analysis of public datasets

Publicly available datasets were obtained from the Gene Expression Omnibus (GEO) database (https://www.ncbi.nlm.nih.gov/geo/) under accession numbers GSE149813, GSE154989, GSE150263, GSE102511, and GSE219124. The study by Dost et al (GSE149813) investigated single lung cells from *Kras^LSL-^*^G12D*;LSL-YFP*^ mice with Ad5CMV-Cre infection ^29^. The report by Marjanovic et al (GSE154989) studied AT2 lineage-labelled cells from lungs of *Kras^LSL-^*^G12D*/+*^*;Rosa26^LSL-tdTomato/+^* mice ^28^. Gene expression count matrices of datasets interrogating *Kras*^G12D^-driven mouse model from GSE149813 were pre-processed using Seurat following the same filtering steps in that original report. For the GSE154989 dataset ^28^, cells used for analysis were the ones labelled as “PASSED_QC” in supplementary table S7 in that study. For the GSE149813 dataset from the Dost et al study, cells with a median number of genes detected > 500 and fraction of mitochondrial genome derived reads < 10%, and according to the pre- processing methods described in their original report ^29^, were retained for analysis. Cells with number of genes detected > 7,500 were further filtered to remove potential doublets/multiplets, resulting in 8,304 cells in total for downstream analysis. Both datasets were integrated with mouse cell data generated in this study using Harmony^18^ with default parameters settings. Top-ranked 20 Harmony corrected PCs were used for clustering with the *FindClusters* function using resolution=0.4. UMAP dimension reduction embedding was performed using the *RunUMAP* function with the same set of Harmony corrected PCs. Gene expression levels and frequencies of representative cluster marker genes were visualized using *DotPlot* function from Seurat. KAC signature score was calculated using the *AddModuleScore* function from Seurat. The mouse KAC signature was also studied in human AT2 cells with and without inducible *KRAS*^G12D^ (dataset GSE150263) also from the study Dost et al ^29^. Cell filtration criteria described in the original report by Dost et al ^29^were followed to filter out potential dead cells and doublets (number of detected genes > 800 and the percent of mitochondrial gene reads fraction < 25%). The 20 top-ranked PCs were used for clustering using the *FindClusters* function with resolution = 0.1. UMAP dimension reduction embeddings were computed using the same SNN graph. The KAC signature score was calculated using *AddModuleScore* function from Seurat package.

The bulk RNA-seq dataset GSE102511 was a previously published dataset by our group comprised of normal lung tissues, precursor atypical adenomatous hyperplasias (AAHs) and matched LUADs (n = 15, each) ^61^. The bulk RNA-seq data GSE219124 by Daouk et al were generated on CSC/stem cell-like progenitor cells, in the form of spheres, and their parental MDA-F471 counterparts (a cell line we had developed and cultured from a KM-LUAD of an NNK-exposed *Gprc5a*^-/-^ mouse) ^62^. To interrogate the association of KACs with tumour formation, gene expression matrices of bulk RNA-seq data GSE102511 (TPM count matrix) and GSE219124 (FPKM count matrix) were extracted and used for quantification of KAC signature expression using MCPcounter (v1.2.0) R package. Heatmaps were generated using pheatmap (v1.0.12) R package.

Mouse KACs from this study were compared to mouse *Krt8*^+^ transitional cells involved in alveolar regeneration post-acute lung injury from the study by Strunz and colleagues^25^. Overlapping marker genes between mouse KACs and the previously reported *Krt8*^+^ transitional cells were statistically evaluated using the ggvenn (v0.1.9) R package using the top-ranked 50 marker genes based on fold change from each study.

### Digital spatial profiling (DSP) of human tissues

The following antibodies were used for DSP: Claudin 4 (Clone 3E2C1, AF594, LSBio, catalogue number LS-C354893, concentration 0.5 µg/ml) and Keratin 8 (Clone EP1628Y, AF647, Abcam, catalogue number ab192468, concentration 0.25 µg/ml). Optimization of antibodies was performed with different dilutions using colorectal carcinoma and LUAD tissues. IF staining was performed on three cases of matched LUAD and NL using the standard GeoMx DSP protocol for morphology markers only (PanCk: clone AE1/AE3, AF532, concentration 0.25 µg/ml, from GeoMx Solid Tumour Morp kit HsP, 121300301, Novus Biologicals, Littleton, CO). Slides were scanned at 20X using the GeoMx DSP platform (NanoString Technologies, Seattle, WA). Following scanning, multiplex IF image slides were visualized adjusting channel thresholds for each fluorophore. Expression of KRT8, PanCK, and CLDN4 was assessed in adenocarcinoma cells, adjacent reactive lung tissue and distant non-reactive lung tissue.

### Animal housing and tobacco carcinogen exposure experiments

Animal experiments were conducted according to Institutional Animal Care and Use Committee (IACUC)–approved protocols at the University of Texas MD Anderson Cancer Center. Mice were maintained in a pathogen-free animal facility. No statistical methods were used to predetermine sample size. In all animal experiments, sex- and age-matched mice were randomized to treatment groups. For all experiments and until endpoints were reached (up to 7 months post-exposure to saline or NNK), mice were monitored for signs of ill health and their body weight was measured to ensure weight loss does not exceed 20% of body weight over 72_Jhours. None of the mice developed these symptoms and thus, they were all euthanized upon reaching IACUC-approved endpoints. Endpoints permitted by our IACUC protocols were not exceeded in any of the experiments. Analysis of data from animal experiments was performed in a blinded fashion. To study KACs in the context of KM-LUAD pathogenesis *in vivo*, *Gprc5a*^−/−^ mice were interrogated since they form LUADs that are accelerated by tobacco-carcinogen exposure and that acquire somatic *Kras*^G12D^ mutations – features that are highly pertinent to KM-LUAD development^21,63,64^ and thus to exploring KACs in this setting. *Gprc5a*^−/−^ mice were generated as previously described^21,65^. Sex- and age-matched *Gprc5a*^−/−^ mice were divided into starting groups of 4 mice per exposure (NNK or saline control) and timepoint (EOE or 7 months post-exposure, n = 16 mice in total). 8-week-old mice were intraperitoneally (IP) injected with 75 mg/kg of body weight nicotine-specific nitrosamine ketone (NNK) or vehicle 0.9% saline (control), three times per week for 8 weeks. At EOE or at 7 months post-exposure, lungs were harvested for derivation of live single cells for scRNA-seq. Whole lungs from additional mice treated as described above were processed by formalin fixation and paraffin-embedding (FFPE) and for analysis by IF (n = 2 mice per treatment group at EOE and 7 months post-exposure, 8 mice in total) and ST (3 lung tissues from *n* = 2 mice at 7 months post-NNK).

*S*ftpc*^CreER/+^; Rosa^Sun1GFP/+^* mice were generously provided by Dr. Harold Chapman (University of California, San Francisco) and were crossed to *Gprc5a^-/-^* mice to generate *Gprc5a^-/-^; S*ftpc*^CreER/+^; Rosa^Sun1GFP/+^* mice for analysis of lineage-labelled AT2 cells. *Gprc5a^-/-^; S*ftpc*^CreER/+^; Rosa^Sun1GFP/+^* mice were treated with 75 mg/kg NNK or control saline (IP), three times per week for 8 weeks. At week 6 of treatment (two weeks before EOE), mice from both groups received 250 µg (IP) tamoxifen dissolved in corn oil for four consecutive days. At EOE or 3 months post-exposure to saline or NNK, lungs were digested to derive live (SYTOX Blue negative) GFP^+^ single cells by flow cytometry using a FACS Aria I instrument as previously described^66^ (gating strategy for GFP cell sorting is shown in **Supplementary Fig. 6**). Sorted single cells were analysed by scRNA-seq (GFP^+^ and GFP^-^ fractions from n = 2 mice per treatment at 3 months post-exposure to saline and NNK) or used to derive organoids (GFP^+^ cells from n = 4 or 5 mice at EOE to saline or NNK, respectively, and from n = 10 or 13 mice at 3 months post-saline or NNK, respectively). Whole lungs from additional mice treated with saline or NNK and tamoxifen as described above (n = 2 per treatment group) were collected (FFPE) at 3 months post-NNK and analysed by IF.

Krt8-CreER; *Rosa^tdT^*^/+^ animals were used to generate *Gprc5a^-/-^;* Krt8-CreER; *Rosa^tdT^*^/+^ mice for analysis of lineage-labelled Krt8^+^ cells. Krt8-CreER (stock number 017947) and *Rosa^tdT^*^/+^ (Ai14; stock number 007914) mice were obtained from the Jackson Laboratory. Mice harbouring Krt8-CreER; *Rosa^tdT^*^/+^ were first used for pilot studies to examine labelling of Krt8^+^ cells. Mice were exposed to control saline (n = 2 mice) or to 8 weeks of NNK (n = 3 mice) like above followed by 1 mg tamoxifen for six continuous days, after which lungs were analyzed at the end of tamoxifen exposure. To examine the relevance of labeled Krt8^+^ cells to tumour development, *Gprc5a^-/-^;* Krt8-CreER; *Rosa^tdT^*^/+^ mice were similarly exposed to NNK for 8 weeks followed by tamoxifen, and lungs were then analyzed at at 8-12 weeks post-NNK (n = 3 mice). All lungs were harvested and processed for formalin fixation, OCT-embedding, and IF analysis.

### Histopathological and IF analysis of mouse lung tissues

Lungs of *Gprc5a^-/-^* mice (n = 2 per treatment and timepoint) were inflated with formalin by gravity drip inflation, excised, examined for lung surface lesions by macroscopic observation, and processed for FFPE, sectioning, and H&E staining. Stained slides were digitally scanned using the Aperio ScanScope Turbo slide scanner (Leica Microsystems Inc., Buffalo Grove, IL) at 200X magnification, and visualized by ImageScope software (Leica Microsystems, Inc.). Unstained lung tissue sections were obtained for IF analysis of Lamp3 (clone 391005, Synaptic Systems), Krt8 (TROMA-I clone from the University of Iowa DSHB), and Pdpn (clone 8.1.1, from the University of Iowa DSHB). Lung FFPE tissue samples were obtained in the same manner from *Gprc5a^-/-^; S*ftpc*^CreER/+^; Rosa^Sun1GFP/+^* mice at 3 months post-exposure to saline or NNK (n = 2 mice per condition) and following injection with tamoxifen. Tissue sections were obtained for H&E staining and assessment of tumour development, and unstained sections were used for IF analysis using antibodies against GFP (AB13970, Abcam, 1:5000), LAMP3 (391005, Synaptic Systems, 1:10000), KRT8 (TROMA-I, University of Iowa Developmental Studies Hybridoma Bank, 1:100), PDPN (clone 8.1.1, University of Iowa Developmental Studies Hybridoma Bank, 1:100), claudin 4 (ZMD.306, Invitrogen, 1:250), and PRKCDBP (Cavin3, Proteintech, 1:250). Slides were then stained with fluorophore-conjugated secondary antibodies and 4’,6’-diamidino-2-phenylindole (DAPI). Sections were mounted with Aquapolymount (18606, Polysciences), cover slipped, imaged using Andor Revolution XDi WD Spinning Disk Confocal microscope and analysed using Imaris software (Oxford Instruments).

Formalin-inflated lung lobes from mice with Krt8-CreER; *Rosa^tdT^*^/+^ were cryoprotected in 20% sucrose in PBS containing 10% optimal cutting temperature compound (OCT; 4583, Tissue-Tek) overnight on a rocker at 4 °C and embedded in OCT the next day, 10 µm cryosections were blocked in PBS with 0.3% Triton X-100 and 5% normal donkey serum (017-000-121, Jackson ImmunoResearch) and incubated overnight in a humidified chamber at 4 °C with primary antibodies diluted in PBS with 0.3% Triton X-100 and raised against: NKX2-1 (sc-13040, Santa Cruz, 1:1000), LAMP3 (same as above) and KRT8 (same as above). The next morning, sections were washed followed by incubation with secondary antibodies (Jackson ImmunoResearch) and DAPI. Slides were then washed, cover slipped as above and imaged using a Nikon A1plus confocal microscope. Cell counter ImageJ plugin was used to count tdT+ cells within lesions, as well as cells in normal-appearing areas, namely: AT2 cells (Lamp3+), tdT+ AT2 cells (tdT+/Lamp3+), AT1 cells (Lamp3-/Nkx2-1+, avoiding noticeable airways), and tdT+ AT1 cells (tdT+/Nkx2-1+/Lamp3-). Percentages of tdT+/Lamp3+ as well as tdT+/Nkx2-1+/Lamp3-cells out of total tdT+ cells were computed. Counts were averages of triplicate images taken at 20X magnification for each time point. The percent regional surface area covered by tdT+ cells in normal-appearing regions was estimated by examining tdT expression across entire lobe sections for each replicate.

### 3D culture and analysis of AT2-derived organoids

*Gprc5a^-/-^;* Sftpc*^CreER/+^; Rosa^Sun1GFP/+^* were treated with NNK or saline as well as tamoxifen as described above, and they were sacrificed at EOE (4 saline- and 5 NNK-treated mice) or at 3 months post-exposure (10 saline- and 13 NNK-treated mice). Lungs were harvested, dissociated into single cells (see mouse single-cell derivation in Methods section “Single-cell isolation from tissue samples”), and live (SYTOX Blue negative) GFP^+^ single cells were collected by flow cytometry using a FACS Aria I instrument as previously described^66^. GFP^+^ AT2 cells from NNK- or saline-treated groups were immediately washed and resuspended at a concentration of 5,000 cells/50 µl of 3D media (F12 medium supplemented with insulin/transferrin/selenium, 10% FBS, penicillin/streptomycin, L-glutamine). GFP^+^ cells were mixed at a 1:1 ratio (by volume) with 50,000 mouse endothelial cells (harvested from mouse lungs by CD31 selection and expanded *in vitro* as previously described ^67^) and resuspended in 50 µl of Geltrex™ reduced growth factor basement membrane matrix (A1413301, Gibco). 100 µl of 1:1 GFP^+^:endothelial cell mixture was plated on transwell inserts with 0.4 µm pores and allowed to solidify for 30 mins in a humidified CO_2_ incubator (EOE: n = 3 wells per condition; 3 months post-exposure: n = 4 wells for saline-derived organoids and n = 12 wells for NNK-derived organoids). Each well was then supplemented with 3D media containing ROCK inhibitor (Y-27632, Millipore) and recombinant mouse FGF-10 (6224-FG, R&D Systems), and plates were incubated at 37°C in a humidified CO_2_ incubator. Wells were replenished with 3D media every other day. For GFP^+^ organoids derived from mice exposed to NNK, 200 nM KRAS^G12D^ specific inhibitor MRTX1133 or DMSO vehicle was added to the media and replenished three times a week (n = 6 wells per condition). Organoids were monitored and analysed twice a week using EVOS M7000 Imaging System (Thermo Fisher Scientific, Waltham, MA), whereby the numbers and sizes of organoids greater than 100 µm in diameter were recorded. At endpoint, 3D organoids were harvested from the basement membrane matrix using Gentle Cell Dissociation Reagent (100-0485, StemCell Technologies), fixed with 4% paraformaldehyde, permeabilized, blocked, and stained overnight at 4°C with a mixture of IF primary antibodies raised against Lamp3, GFP, Krt8, and Cavin3. The next day, organoids were washed and stained with fluorophore-conjugated secondary antibodies overnight at 4°C while being protected from light. Organoids were washed and stained with DAPI nuclear stain for 30 minutes, after which they were collected in Aqua-Poly/Mount (18606-20, Polysciences) and transferred to slides. Images of organoids were captured using an Andor Revolution XDi WD Spinning Disk Confocal microscope and analysed using Imaris software (Oxford Instruments, Abingdon, United Kingdom).

### 2D viability assays

Mouse mycoplasma-free LUAD cell lines LKR13 (mutant *Kras*^G12D^-driven; ^31^) and MDA-F471 (*Gprc5a*^-/-^ and *Kras*^G12D^ mutant; ^27^) were plated on 96-well plates (10^3^ cells/well) and grown in DMEM (Dulbecco’s Modified Eagle Medium, Gibco), supplemented with 10% FBS, 1% antibiotic antimycotic solution (A5955, Sigma-Aldrich, St. Louis, MO), and 1% L-Glutamine (G7513, Sigma-Aldrich, St. Louis, MO). The next day, cells were cultured for up to 4 days with media containing either 0.5% FBS, 0.5% FBS with 50 ng/ml epidermal growth factor (EGF) (E5160, Sigma-Aldrich, St. Louis, MO), or 0.5% FBS with EGF and varying concentrations of MRTX1133 (Mirati Therapeutics, San Diego, CA). 25 ul alamarBlue Cell Viability Reagent (DAL1025, ThermoFisher, Waltham, MA) was added to each well. At 4 days post-treatment, viability was assessed by fluorescence spectrophotometry at 570 nm (and 600 nm as a reference) where for wells showing net positive absorbances relative to blank wells (at least 3 wells per cell line and condition) percent differences in reduction between treated and control wells were calculated.

### Western blot analysis

LKR13 and MDA-F471 cells were plated in 6-well plates (10^6^ cells/well) and grown under different conditions as described above. Protein lysates were extracted at 3 hours post-treatment and analysed by western blot following overnight incubation with antibodies against the following primary proteins: Vinculin (E1E9V, rabbit, Cell Signaling Technology, 13901, 1:1000), Phosphorylated p44/42 MAPK (ERK1/2, rabbit, Cell Signaling Technology, 9101, 1:2000), Phosphorylated S6 Ribosomal protein (Ser 235/236, rabbit, Cell Signaling Technology, 4858, 1:2000), p44/42 MAPK (ERK1/2, rabbit, Cell Signaling Technology, 9102, 1:2000), or S6 (E.573.4, rabbit, Invitrogen, MA5-15164, 1:1000), and followed by one-hour incubation with diluted secondary antibody (1706515 Goat Anti-Rabbit IgG-HRP Conjugate, Bio-Rad, Hercules, CA). Protein lysates from each cell line were analysed on multiple gels (four per cell line) with Precision Plus Protein Dual Color Standard (1610394, Bio-Rad, Hercules, CA) as ladder and blotted to membranes to separately probe for phosphorylated and total forms of the same proteins, which have very similar molecular weights (using phospho-specific antibodies or antibodies targeting total version of same protein). Vinculin protein levels were evaluated as loading control on each of the blots. Four blots (phospho-ERK, total ERK, phospho-S6, total S6) for each of LKR13 and MDA-F471 are shown in **Supplementary Fig. 9, each with its own analysis of equal protein loading** (vinculin blot) and whereby only the ones indicated with green rectangles are presented in **Extended Data Fig. 12c**. Membranes were cut horizontally using molecular weight marker as a guide, and cut membranes were incubated with the specified antibodies (see **Supplementary Fig. 9 for site of cutting and for overlay of colorimetric and chemiluminescent images of the same blot to display ladder and the analysed protein, respectively**). Blots were imaged using the ChemiDoc Touch Imaging System (Bio-Rad, Hercules, CA) with “Chemiluminescence” and “Colorimetric” (for protein ladder) applications and auto expose or manual settings.

### Chemicals and reagents

Tobacco-specific carcinogen (nicotine-specific nitrosamine ketone; NNK) with a purity of 99.96% by HPLC was purchased from TargetMol (Wellesley Hills, MA). Tamoxifen, as well as H&E staining reagents, were purchased from Sigma Aldrich (St. Louis, MO). The KRAS^G12D^ inhibitor MRTX1133 was generously provided by Dr. James Christensen (Mirati Therapeutics Inc., San Diego, CA).

### Statistical analyses

In addition to the algorithms and statistical analyses described above, all other basic statistical analyses were performed in the R statistical environment (version 4.0.0). The Kruskal–Wallis H test was used to compare variables of interests across three or more groups. Wilcoxon Rank-Sum test was used for paired comparisons among matched samples from the same patients. Wilcoxon Rank-Sum test was used to compare other continuous variables such as gene expression levels and signature scores between groups. Spearman’s correlation coefficient was calculated to assess associations between two continuous variables (e.g., cellular proportions and gene signature scores). Fisher’s exact test was used to identify differences in frequencies of groups based on two categorical variables. Ordinal logistic regression was performed using the *polr* function in the built-in R package MASS (v7.3). Benjamin-Hochberg method was used to control for multiple hypothesis testing. All statistical tests performed in this study were two-sided. Results were considered statistically significant at *P* - values or FDR *q* - values < 0.05. When a *P* - value reported by R was smaller than 2.2e-16, it was reported as “*P* < 2.2 × 10^-16^”.

## Data Availability

Sequencing data for P1 - P5 were previously generated and deposited in the European Genome– phenome Archive (EGA) under the accession number EGAS00001005021^15^. Human scRNA-seq (P6 – P16) and ST data generated in this study are deposited in EGA under the same accession number (EGAS00001005021). Mouse scRNA-seq and ST data generated in this study are deposited in NCBI under GEO accession number GSE222901. Relevant source data are provided with this paper.

## Code availability

Codes for analysis of scRNA-seq, WES, and ST data are available at Zenodo (https://doi.org/10.5281/zenodo.8280138) and GitHub (https://github.com/guangchunhan/LUAD_Code).

## Acknowledgements

We appreciate all the patients who generously provided their samples for this research under approved consents. We thank Dr. Harold Chapman for kindly providing *S*ftpc*^CreER/+^; Rosa^Sun1GFP/+^* mice. This work was supported in part by funding from Johnson and Johnson (to H.K.), The University of Texas MD Anderson Cancer Center Office of Strategic Research Programs (to H.K.), the National Cancer Institute (NCI) grants R01CA205608 (to H.K.), R01CA272863 (to H.K.), 1U2CCA233238 (to H.K., S.M.D., and A.E.S.), U01CA264583 (to H.K. and L.W.) and T32CA217789 MD Anderson Cancer Center postdoctoral fellowship (to A.S.), University of Texas SPORE in Lung Cancer P50CA070907, as well as the Cancer Prevention and Research Institute of Texas grants RP150079 (to H.K.) and RP220101 (to H.K.). L.W. and H.K. acknowledge support from the start-up research fund from the University of Texas MD Anderson Cancer Center. T.C., L.W. and H.K. are Andrew Sabin Family Foundation Fellows at the University of Texas MD Anderson Cancer Center. The Flow Cytometry & Cellular Imaging Core Facility is supported by the National Cancer Institute through MD Anderson’s Cancer Center Support Grant P30 CA016672. The sequencing core facility at MD Anderson Cancer Center is supported by the National Institutes of Health grant 1S10OD024977.

## Author information

These authors contributed equally: Guangchun Han, Ansam Sinjab These authors contributed equally: Zahraa Rahal, Anne M. Lynch These authors jointly supervised the work: Jichao Chen, Linghua Wang, Humam Kadara

### Authors and Affiliations

**Department of Translational Molecular Pathology, The University of Texas MD Anderson Cancer Center, Houston, TX, USA**

Ansam Sinjab, Zahraa Rahal, Warapen Treekitkarnmongkol, Yuejiang Liu, Alejandra G. Serrano, Jiping Feng, Ke Liang, Khaja Khan, Wei Lu, Sharia Hernandez, Camille Abaya, Lorena I. Gomez-Bolanos, Minyue Chen, Edwin R. Parra, Junya Fujimoto, Luisa M. Solis, Ignacio I. Wistuba & Humam Kadara

**Department of Genomic Medicine, The University of Texas MD Anderson Cancer Center, Houston, TX, USA**

Guangchun Han, Yunhe Liu, Xuanye Cao, Enyu Dai, Guangsheng Pei, Fuduan Peng & Linghua Wang

**Department of Pulmonary Medicine, The University of Texas MD Anderson Cancer Center, Houston, TX, USA**

Anne M. Lynch, Seyed Javad Moghaddam & Jichao Chen

**Graduate Program in Developmental Biology, Baylor College of Medicine, Houston, Texas, USA**

Anne M. Lynch

**Department of Thoracic/Head and Neck Medical Oncology, The University of Texas MD Anderson Cancer Center, Houston, TX, USA**

Tina Cascone, Marcelo V. Negrao & John V. Heymach

**Department of Cardiovascular and Thoracic Surgery, The University of Texas MD Anderson Cancer Center, Houston, TX, USA**

Boris Sepesi

**Department of Epidemiology, The University of Texas MD Anderson Cancer Center, Houston, TX, USA**

Paul Scheet

**Department of Biostatistics, Epidemiology and Informatics, Perelman School of Medicine, University of Pennsylvania, Philadelphia, PA, USA**

Jian Hu & Mingyao Li

**Department of Medicine, The University of California Los Angeles, Los Angeles, CA, USA**

Steven M. Dubinett

**Lung Cancer Initiative at Johnson and Johnson, Boston, MA, USA**

Christopher S. Stevenson & Avrum E. Spira

**Section of Computational Biomedicine, School of Medicine, Boston University, Boston, MA, USA**

Avrum E. Spira

**The University of Texas Health Houston Graduate School of Biomedical Sciences, Houston, TX, USA**

Yuejiang Liu, Minyue Chen, Tina Cascone, Seyed Javad Moghaddam, Paul Scheet, John V. Heymach, Linghua Wang & Humam Kadara

### Contributions

G.H., A.S., L.W. and H.K. designed the study, interpreted the data, and wrote the original draft of the manuscript. All authors reviewed the final version of the manuscript. G.H. led computational analyses relating to scRNA-seq, WES and ST. G.H., A.S., L.W. and H.K. performed data quality control and curation. G.H., X.C. and F.P. processed and aligned scRNA-seq and ST data. G.H. and E.D. analysed MPs. G.H. and Yunhe.L. performed *KRAS* mutation screening using the scRNA-seq data. G.H., Yunhe.L. and G.P. curated bioinformatic pipelines for ST analysis. G.H., A.S., L.W. and H.K. annotated cells, governed overall analysis and interpretation of scRNA-seq, ST and WES data as well as performed data visualization. P.S. developed workflows for analysis of mutations in normal tissues and assisted in data interpretation. A.S. led generation of scRNA-seq, WES, ST and IF data. A.S., K.K. and L.M.S. processed tissues for ST analysis. J. Fujimoto performed histopathological analysis of tissues analysed by scRNA-seq. A.G.S., J. Fujimoto and L.M.S. performed spot-level histopathological evaluation of mouse and human tissues analysed by ST. W.L., S.D.H. and L.M.S. performed the digital spatial profiling of human tissues. L.I.B-G., E.R.P., L.M.S. developed workflows and provided resources for digital spatial profiling and IF analysis. A.S., Z.R., J. Feng and W.T. performed tobacco carcinogenesis experiments. A.S. and Z.R. performed tobacco carcinogenesis experiments in AT2 and Krt8 reporter mice. A.S., Z.R., A.M.L., K.L., J.C. and H.K. analysed AT2 and KRT8 tracing in reporter-containing tobacco carcinogenesis experiments. A.M.L., S.J.M. and J.C. developed workflows for Krt8 reporter tracing and IF analysis of mouse lung tissues. J.H. and M.L. developed workflows and tools for ST analysis used in this study. M.L. assisted in human ST data interpretation and analysis. A.S., Yuejiang.L., C.A. and M.C. performed experiments with cell lines and organoids. T.C., B.S., M.V.N., J.V.H. and J. Fujimoto provided human tissue resources and clinical annotations. M.V.N. and J.V.H. provided advice for KRAS targeted inhibition studies. S.M.D., C.S.S. and A.E.S. provided administrative support, data interpretation and resources pertaining to lung cancer cohorts. J.C., L.W. and H.K. supervised the overall study. H.K. provided strategic oversight and conceived the study.

### Corresponding author

Correspondence to Jichao Chen, Linghua Wang, or Humam Kadara.

## Ethics declarations

All human LUAD and normal lung tissues were obtained from patients who provided informed consents and under IRB-approved protocols at The University of Texas MD Anderson Cancer Center. All human data in this manuscript are deidentified to ensure patient privacy. All animal studies were conducted under IACUC-approved protocols at the University of Texas MD Anderson Cancer Center.

## Additional information

Supplementary Figs. 1-9 and Supplementary Tables 1-12 are included in Supplementary Information accompanying this manuscript.

## Extended Data Figure Legends

**Extended Data Fig. 1. Analysis of normal lung epithelial and malignant subsets in early-stage LUADs. a, b,** UMAP plots of 229,038 normal epithelial cells from 63 samples. Each dot represents a single cell coloured by major cell lineage (**a**, left), airway sub-lineage (**a**, top right) and alveolar sub-lineages (**a**, bottom right). SDP cells were separately coloured to show their position on the UMAP (**b**). **c, d** UMAP plots of 17,064 malignant cells coloured by patient ID (**c**, left), CNV score (**c**, middle), presence of *KRAS*^G12D^ mutation (**c**, right) and smoking status (**d**). **e,** Analysis of recurrent driver mutations identified by WES. **f,** Transcriptomic variances quantified by Bhattacharyya distances at the sample (left) and cell (right) levels among LUADs with driver mutations in *KRAS* (KM), *EGFR* (EM), and *MET* (MM), or LUADs that are wild type (WT) for these genes. Box, median ± interquartile range; whiskers, 1.5× interquartile range; centre line: median. *n* cells in each box-and-whisker in the left panel: KM-KM = 3; KM-EM = 15; KM-MM = 6; KM-Other = 12; EM-EM = 10; EM-MM = 10; EM-Other = 20; MM-Other = 8; Other-Other = 6. *n* cells in each box-and-whisker in the right panel: 100. *P* - values were calculated by two-sided Wilcoxon Rank-Sum test with a Benjamini–Hochberg correction. **g,** Harmony-corrected UMAP plot of malignant cells coloured by cluster ID (left) and cluster distribution by sample (right). **h,** UMAP plots of malignant cells coloured by CNV scores (top left), smoking status (top right). Comparison of CNV scores between malignant cells from samples carrying different driver mutations (bottom left) or between smokers and never smokers (bottom right). Box-and-whisker definitions are similar to panel **f**. *n* cells in each box-and-whisker: *EGFR* = 5,457; Other = 9,135; *KRAS* = 2,472; Smoker = 5,999; Never smoker = 11,065. *P* - values were calculated by two-sided Wilcoxon Rank-Sum test with a Benjamini–Hochberg correction. **i,** Analysis of Wasserstein distances among KM-LUADs, EM-LUADs, and LUADs with WT *KRAS* and *EGFR* (Double WT). Box-and-whisker definitions are similar to panel **f**. *n* samples in each box-and-whisker: 3; 5; 6. *P* - value was calculated by a two-sided Wilcoxon Rank-Sum test.

**Extended Data Fig. 2. Characterization of inter- and intra-tumour heterogeneity of LUAD malignant cells. a,** Unsupervised clustering of malignant cells based on expression of 23 previously defined consensus cancer cell meta-programs (MPs). **b**, Distribution of signature scores of 4 representative MPs across clusters from **a**. Box-and-whisker definitions similar to **Extended Data Fig. 1f**. *n* cells in each box-and-whisker: C1 = 2,600; C2 = 3,968; C3 = 1,647; C4 = 7,182; C5 = 1,667. **c**, Enrichment of clusters (C1-C5) in cells colour coded by recurrent driver mutation status (left) and patients (right). **: *P* < 2.2 × 10^-16^. *P* - value was calculated using two-sided Fisher’s exact test with a Benjamini–Hochberg correction. **d,** MP30 was computed in malignant cells in each patient (left) and in KM-LUADs versus *KRAS* WT LUADs (KW-LUADs, right). *n* cells in each box-and-whisker: P14 = 1,614; P10 = 326; P2 = 532; P1 = 64; P6 = 2,604; P7 = 823; P8 = 147; P15 = 1,819; P4 = 404; P9 = 25; P3 = 2,419; P5 = 5,872; P11 = 375; P13 = 40; KM-LUADs = 2,472; KW-LUADs = 14,592. Box-and-whisker definitions are similar to **Extended Data Fig. 1f**. *P* - values were calculated using two-sided Wilcoxon Rank-Sum test with a Benjamini–Hochberg correction. **e**. Profiling of ITH in malignant cells from P14 LUAD. UMAP plots show malignant cells coloured by (top left to top right) *KRAS*^G12D^ mutation status, KRAS signature expression, and cell differentiation status (CytoTRACE). Trajectories of P14 malignant cells coloured by (bottom left to bottom right) the presence of *KRAS*^G12D^ mutation, inferred pseudotime, and differentiation status. **f,** UMAP plots showing P14 malignant cells coloured by expression of the 3 indicated MPs. **g**, Unsupervised clustering analysis of P14 malignant cells based on inferred CNV profiles (left). UMAP of P14 malignant cells (middle) and inferred trajectory (top right) coloured by CNV clusters, as well as *KRAS*^G12D^ mutation expression status along pseudotime trajectory (bottom right). **h**, Alveolar MP expression across the CNV clusters shown in panel **g**. *n* cells in each group: 477; 464; 673*. P* - values were calculated using two-sided Wilcoxon Rank-Sum test with a Benjamini–Hochberg correction. **i**, Harmony-corrected UMAP plot of malignant cells coloured by KRAS signature score (left). Correlation between MP30 expression and KRAS signature score in malignant cells of KM-LUADs (right). *P* - value was calculated with Spearman correlation test. R denotes the Spearman correlation coefficient. **j**, Heatmap showing score distribution of the indicated MPs and signatures in TCGA LUAD samples. **k,** Kaplan-Meier plot showing differences in the survival probability between samples with high and low levels of KRAS signature (KRAS sig.), and those with *KRAS*^G12D^ mutation. OS: overall survival. *KRAS* sig. high: samples within top quartile of KRAS signature score. *KRAS* sig. low: samples below the third quartile of KRAS signature score. mo.: months. *P* - value was calculated with logrank test.

**Extended Data Fig. 3. Phenotypic diversity and states of human normal lung epithelial cells. a,** Composition of normal epithelial lineages across spatial regions as defined in **Fig. 1a**. Dis: distant normal. Int: intermediate normal. Adj: adjacent normal. NE: neuroendocrine. **b,** Changes in cellular fractions of AT2 cells (left) and AICs (right) across the spatial samples. Box-and-whisker definitions are similar to **Extended Data Fig. 1f**. *n* samples in each box-and-whisker (left to right): 16; 15; 16; 16. *P* - values were calculated with Kruskal-Wallis test. **c,** Composition of normal epithelial lineages across the spatial regions at the sample level. **d**, Fractional changes of AT2 cells among all epithelial cells across the spatial regions at the patient level. **c** and **d**: Cases showing gradually reduced AT2 fractions with increasing tumour proximity (7 of the 16 patients; *P* = 0.004 by ordinal regression analysis in **d**). **e,** Fractions of AT1, basal, ciliated, and club and secretory cells along the continuum of the spatial samples. Box-and-whisker definitions are similar to **Extended Data Fig. 1f**. *n* samples in each box-and-whisker (left to right): 16; 16; 15; 16. *P* - values were calculated with Kruskal-Wallis test. **f**, Distribution of CytoTRACE scores in AICs, AT1 and AT2 cells (left). Distribution of pseudotime scores in malignant cells from *EGFR*- or *KRAS*-mutant tumours (right). *P* - value was calculated with two-sided Wilcoxon Rank-Sum test. Box-and-whisker definitions are similar to **Extended Data Fig. 1f** with *n* cells: AT2 = 14,649; AICs = 974; AT1 = 2,529; *EGFR* = 1,711; *KRAS* = 1,326. **g**, Pseudotime trajectory analysis of alveolar and malignant subsets coloured by tissue location. **h**, Distribution and composition of AICs with low (left) or high (right) CytoTRACE score. **i**, DEGs between KACs and other AICs. **j,** Pseudotime trajectory analysis of malignant and alveolar subsets colour-coded by cell lineage and presence of *KRAS*^G12D^ mutation (top). Pseudotime score in KACs versus other AICs (bottom). Box-and-whisker definitions are similar to **Extended Data Fig. 1f**. *n* cells in each box-and-whisker: KACs = 157; Other AICs = 817. *P -* value was calculated by two-sided Wilcoxon Rank-Sum test. **k**, Differences in cell densities between LUAD (top) and NL tissues (bottom).

**Extended Data Fig. 4. Spatial and molecular attributes of human KACs. a,** Microphotographs of P10 (left) and P15 (right) LUAD and paired uninvolved NL tissues. Top panels: H&E staining showing LUAD T and TAN (left columns) regions, and uninvolved NL (right columns). DSP analysis of KRT8 (red), CLDN4 (yellow), and pan-cytokeratin (PanCK; green) in LUAD, TAN, and NL regions. Blue nuclear staining was done using Syto13. Magnification 20X. Scale bar = 200 μm. Staining was repeated four times with similar results. **b,** CytoSPACE deconvolution and trajectory analysis of P14 LUAD ST data. The left spatial map is coloured by deconvoluted cell types. Top middle panel shows the neighbouring cell composition of KACs, and the bottom middle panel depicts inferred trajectory and pseudotime prediction using Monocle 2. Scaled expression of *NKX2-1* and alveolar signature are shown in the rightmost top and bottom panels, respectively. **c-e,** Expression of KRAS (**c**), AT1 (**d**), and other AIC (**e**) signatures across AT1, AT2, KACs and other AICs. Box-and-whisker definitions are similar to **Extended Data Fig. 1f**. *n* cells in each group: KACs = 1,440; Other AICs = 8,593; AT2 = 146,776; AT1= 25,561. **f, g,** Correlation analysis between Other AIC and KRAS (**f**) or alveolar (**g**) signature scores. *P* - values were calculated with Spearman correlation test. R denotes the Spearman correlation coefficients. **h,** Enrichment of KAC signature among KACs (left) and malignant cells (right) from KM- or EM-LUAD samples. Box-and-whisker definitions are similar to **Extended Data Fig. 1f**. *n* cells in each box-and-whisker (left to right): KACs, EM- LUADs = 135; KACs, KM-LUADs = 719; Malignant, EM-LUADs = 5,457; Malignant, KM- LUADs = 2,472. *P -* values were calculated by two-sided Wilcoxon Rank-Sum test.

**Extended Data Fig. 5. Enrichment and clinical relevance of KAC, Other AIC, and alveolar signatures in LUAD. a-e,** Expression of KAC (**a**), other AIC (**b**) and alveolar (**c**) signatures in TCGA LUAD samples and matched NL tissues, of other AIC signature in a lung preneoplasia cohort (**d**), as well as of KAC signature in TCGA LUAD samples grouped by *KRAS* mutation status (**e**). Box-and-whisker definitions are similar to **Extended Data Fig. 1f**. *n* samples in each group: TCGA Normal = 52; TCGA LUAD = 52; preneoplasia Normal, AAH, and LUAD: 15 each; TCGA LUAD *KRAS* WT = 346; TCGA LUAD *KRAS* MUT = 152. *P -* values were calculated by two-sided Wilcoxon Rank-Sum test. Benjamini–Hochberg method was used for multiple testing correction. n.s.: non-significant (*P* > 0.05). **f-i,** Kaplan-Meier plots showing differences in overall survival probability across TCGA (**f**) and PROSPECT (**g**) samples with high versus low KAC signature scores, or with high versus low scores for other AIC signature (**h**: TCGA; **i**: PROSPECT). Sig. low: LUAD samples with signature scores lower than the group median value. Sig. hi: LUAD samples with signature scores higher than the group median value. *P* - values were calculated with the logrank test. **j,** Multivariate Cox proportional hazard regression analysis including pathologic stage, age, as well as KAC and other AIC signatures. Center: estimated Hazard Ratio; error bars: 95% CI. *q* - values were calculated by Cox proportional hazards regression model and adjusted with Benjamini–Hochberg method.

**Extended Data Fig. 6. Prevalence of *KRAS*^G12D^ mutant KACs in LUAD. a,** UMAP clustering of alveolar subsets. **b**, Quantification of CNV scores across AT1, AT2, KACs and other AICs. Box-and-whisker definitions are similar to **Extended Data Fig. 1f**. *n* cells in each group: AT2 =146,776; AT1 = 25,561, Other AICs =8,593; KACs = 1,440; Malignant = 17,064. *P* – values were calculated using two-sided Wilcoxon Rank-Sum test with a Benjamini–Hochberg correction. *KRAS*^G12D^ variant allele frequencies (**c**) and fractions of *KRAS*^G12D^ mutant cells (**d**) in alveolar and malignant cells from LUAD and normal samples and analysed by scRNA-seq. VAF for *KRAS*^G12C^ variant in KACs from KM normal tissues is shown in green (**c**). *n* on top of each bar in **d**: number of *KRAS*^G12D^ mutant cells. **e**, KRAS activation signature was statistically compared across *KRAS*^G12D^ mutant KACs, *KRAS*^wt^ KACs, AICs, and AT2 cells. Box-and- whisker definitions are similar to **Extended Data Fig. 1f**. *n* cells in each box-and-whisker: KACs *KRAS*^G12D^ = 15; KACs *KRAS*^wt^ = 1,425; Other AICs =8,593; AT2 = 146,776. *P* – values were calculated using the two-sided Wilcoxon Rank-Sum test with a Benjamini–Hochberg correction. **f, g,** CytoTRACE scores in KACs versus other AICs from all cells of KM (**f**, left) and KW cases (**f**, right), in cells from normal lung tissues of patients with KM-LUAD (**g**, left), and cells from KM-LUAD (**g**, middle) and KW-LUAD (**g**, right) tissues. Box-and-whisker definitions are similar to **Extended Data Fig. 1f**. *n* cells in each box-and-whisker: KM cases, KACs = 719; KM cases, Other AICs = 2,414; KW cases, KACs = 721; KM cases, Other AICs = 6,179; KM normal tissues, KACs = 408; KM normal tissues, Other AICs = 2,286; KM-LUADs, KACs = 311; KM-LUADs, Other AICs = 128; KW-LUADs, KACs = 295; KW-LUADs, Other AICs = 940. *P* - values were calculated using two-sided Wilcoxon Rank-Sum tests with Benjamini–Hochberg adjustment for multiple testing correction.

**Extended Data Fig. 7. scRNA-seq analysis of epithelial subsets in a tobacco carcinogenesis mouse model of KM-LUAD. a,** UMAP distribution of mouse epithelial cell subsets. **b,** Proportions and average expression levels of select marker genes for mouse normal epithelial cell lineages and malignant cell clusters as defined in panel **a**. **c,** UMAP plots of alveolar and malignant cells coloured by CNV score, presence of *Kras*^G12D^ mutation, or expression levels of *Kng2* and *Meg3*. **d,** UMAP (top) and violin (bottom) plots showing expression level of *Cd24a* in malignant and alveolar subsets. Box-and-whisker definitions are similar to **Extended Data Fig. 1f**. *n* cells in each group: Malignant = 1,693; AT1 = 580; KACs = 636; AT2 = 1,791. **e**, UMAP distribution of alveolar and malignant cells coloured by cell lineage, *Kras*^G12D^ mutation status, and CNV score at EOE or 7 mo post-NNK. **f**, Proportions of normal epithelial cell lineages and malignant cells in each sample. **g**, Fractional changes of malignant cells, KACs, AT2 and AT1 cells between EOE and 7 months post treatment with NNK or saline; n = 4 biologically independent samples in each group. Whiskers, 1.5× interquartile range; Center dot: median. **h**, UMAP (top) and violin (bottom) plots showing expression levels of *Gkn2* in malignant and alveolar cell subsets. *n* cells in each group: Malignant = 1,693; AT1 = 580; KACs = 636; AT2 = 1,791.

**Extended Data Fig. 8. ST analysis of KACs in tobacco-associated development of KM- LUAD. a,** ST analysis of the same tumour-bearing mouse lung in **Fig. 3e** with cell clusters identified by Seurat (inlet) and mapped spatially (left). Spatial maps with scaled expression of *Krt8* and *Plaur* are shown on the right. **b**, Pseudotime trajectory analysis of C0 (alveolar parenchyma), C2 (reactive area with KACs nearby tumours), and clusters C7 and C8 (representing two tumours) from the same tumour-bearing mouse lung in **a**. **c**, ST analysis of another tumour-bearing lung region from the same NNK-exposed mouse as in panel a, and showing histological spot-level annotation of H&E-stained images (left) followed by spatial maps with scaled expression of *Krt8*, *Plaur*, and KAC signature (right). **d**, Cell clusters identified by Seurat (top left) and mapped spatially (top right) from the same mouse tumour-bearing lung in **c**. bottom of panel k: Pseudotime trajectory analysis of C0 (alveolar parenchyma), C8 (reactive area with KACs nearby the tumour), and C5 (representing one tumour) from the mouse tumour-bearing lung in **c**. **e**, ST analysis of a tumour-bearing lung from an additional mouse at 7 months following NNK showing histological spot-level annotation of H&E-stained images (left) followed by spatial maps with scaled expression of *Krt8* (middle, top), *Plaur* (middle, bottom), and KAC signature (right).

**Extended Data Fig. 9. Mouse KAC signatures and pathways are relevant to both injury models and human KM-LUAD. a, b,** Pathway enrichment analysis of KACs relative to other alveolar cell subsets and malignant cells in tumour-bearing mice at 7 months post-NNK (**a**) and in the human LUAD scRNA-seq dataset from this study (**b**). **c,** Enrichment of *Tp53* signature derived from mouse KACs, and expression of *Btg2*, *Ccng1*, *Cdkn2b*, *Bax*, *Cdkn1a*, as well as *Trp53* itself, across AT2 cells, malignant cells, and KACs at EOE or at 7 mo post-NNK or saline. *n* cells in each group: AT2 = 1,791; KACs EOE = 301; KACs 7mo. = 335; Malignant =1,693. **d**, Pie chart showing percentages of unique and overlapping DEG sets between mouse KACs from this study and *Krt8^+^* transitional cells identified by Strunz and colleagues. **e, f,** Expression of the mouse KAC signature across alveolar and malignant cell subsets from this study (**e**), in normal lung (Normal) and LUAD tissues from the TCGA cohort (**f,** left), as well as in normal lung (Normal), AAH, and LUAD tissues of our premalignancy cohort (**f,** right). *n* cells in each group of panel **e**: AT2 = 1,791; KACs EOE = 301; KACs 7mo. = 335; Malignant = 1,693. *n* samples in each group of panel **f** left: Normal = 52; LUAD = 52. *n* samples in each group of panel **f** right: Normal = 15; AAH = 15; LUAD = 15. Box-and-whisker definitions are similar to **Extended Data Fig. 1f**. *P* - values were calculated using two-sided Wilcoxon Rank-Sum test with a Benjamini–Hochberg correction.

Extended Data Fig. 10. Mouse KACs exist in a continuum, bear strong resemblance to human KACs, and are present in independent *KRAS*^G12D^-driven mouse models of LUAD. a, Mouse KAC signature score (left) and heatmap showing expression of select KAC marker genes (right) in bulk transcriptomes of MDA-F471-derived 3D spheres versus parental MDA-F471 cells grown in 2D. *P* - value was calculated using two-sided Wilcoxon Rank-Sum test. Box-and-whisker definitions are similar to **Extended Data Fig. 1f**. **b**, Fraction of *Kras*^G12D^ mutant cells in different mouse alveolar cell subsets including when separating KACs into early KACs at EOE and late KACs at 7 months post-NNK. Numbers of *Kras*^G12D^ mutant cells are indicated on top of each bar. **c,** CytoTRACE scores in late KACs with *Kras*^G12D^ mutation and in those with wild type *KRAS* (*Kras*^wt^). *P* - value was calculated using two-sided Wilcoxon Rank-Sum test. Box-and-whisker definitions are similar to **Extended Data Fig. 1f**. *n* cells in each box-and-whisker: *Kras*^G12D^ = 72; *Kras*^wt^ = 564. **d,** Proportions and average expression levels of select marker genes for the different subsets indicated. Pie charts showing percentages of unique and overlapping DEG sets between *Krt8^+^* transitional cells identified by Strunz and colleagues and either *Kras*^G12D^ (**e**) or *Kras*^wt^ (**f**) KACs from this study. **g,** UMAP clustering of cells integrated from our mouse cohort with cells in the scRNA-seq datasets from studies by Marjanovic et al. and Dost et al. **h,** Proportions and average expression levels of select marker genes for diverse alveolar and tumour cell subsets and across clusters defined in panel **g** with cluster 5 (C5) shown to be enriched with KAC markers. **i,** KAC signature expression across clusters defined in panel g. *n* cells in each cluster: 2 = 2,463; 11 = 154; 1 = 3,480; 0 = 4,396; 5 = 1,362; 4 = 1,513; 3 = 2,392; 10 = 219; 8 = 577; 7-0 = 382; 6 = 1,042; 9 = 285; 7-1 = 141; 7-2 = 115; 12 = 119. **j,** Distribution of cells from C5 across the three indicated cohorts (left). KAC signature enrichment across KACs from the three cohorts and relative to pooled AT2 cells (right). Box-and-whisker definitions are similar to **Extended Data Fig. 1f**. *n* cells in each box-and-whisker: KACs, Marjanovic et al = 90; This study = 485; Dost et al = 343; AT2 = 3,762. **k,** KAC signature score in human AT2 cells with induced expression of *KRAS*^G12D^ (Dox) relative to *KRAS*^wt^ cells (Ctrl) from the Dost et al. study. Dox: Doxycycline. Box-and-whisker definitions are similar to **Extended Data Fig. 1f**. *n* cells in each box-and-whisker: Ctrl = 802; Dox = 1,341. *P* - value was calculated using two-sided Wilcoxon Rank-Sum test**. l,** Mouse KAC signature expression in KACs (left) and malignant cells (Malignant, right) from KM-LUADs relative to EM-LUADs in our human scRNA-seq dataset. Box-and-whisker definitions are similar to **Extended Data Fig. 1f**. *n* cells in each box-and-whisker: KACs, EM-LUADs = 135; KACs, KM-LUADs = 719; Malignant, EM-LUADs = 5,457; Malignant, KM-LUADs = 2,472. *P* - values were calculated using two-sided Wilcoxon Rank-Sum test.

Extended Data Fig. 11. KACs are enriched in lungs and they precede the formation of *Kras*^G12D^ tumours in an AT2 lineage reporter tobacco carcinogenesis mouse model. a, Representative IF analysis of KRT8, GFP, and LAMP3 in GFP-labelled AT2-derived mouse lung organoids (n = 3 wells per condition) derived from tamoxifen-exposed AT2 reporter mice at EOE to saline (n = 4 mice) or NNK (n = 5 mice). Scale bar: 10 μm. **b**, UMAP distribution of GFP^+^ cells at 3 months following NNK exposure or saline and coloured by alveolar or tumour subsets. **c**, Proportions and average expression levels of select marker genes for mouse normal alveolar cell lineages and tumour cells defined in **b**. **d,** Fraction of *Kras*^G12D^ cells across alveolar and early tumour subsets. Absolute numbers of *Kras*^G12D^ cells are indicated on top of each bar**. e,** UMAPs of GFP^+^ cells from tumour-bearing AT2 reporter mice at 3 months post-NNK or saline and coloured by presence of *Kras*^G12D^ mutation or expression of KAC, AT1, and AT2 signatures. **c,** UMAPs showing distribution of alveolar and tumour cell subsets (left) as well as cells with *Kras*^G12D^ mutation (right) by treatment (saline or NNK). **f,** Trajectories of GFP^+^ cells from tumour-bearing reporter mice at 3 months post-NNK or saline coloured by inferred pseudotime (left), differentiation (middle), and cell lineage and showing subset composition (right). **g,** CytoTRACE (left) and pseudotime (right) scores across GFP^+^ subsets. Box-and-whisker definitions are similar to **Extended Data Fig. 1f**. *n* cells in each box-and-whisker: AT2 = 144; Early/AT2-like tumour = 144; KAC/KAC-like = 288; AT1 = 72.

**Extended Data Fig. 12. KAC-rich organoids are sensitive to targeted inhibition of KRAS. a,** Size quantification of organoids derived from GFP^+^ lungs cells of mice treated with saline (derived from 10 mice and plated into 4 wells) or NNK (derived from 13 mice and plated into 12 wells) at 3 months post-exposure. Box-and-whisker definitions are similar to **Extended Data Fig. 1f**. *n* organoids in each group: Saline = 63; NNK = 66. *P* - value was calculated using two-sided Wilcoxon Rank-Sum test. **b,** Analysis of relative viability 4 days post treatment of LKR13 and MDA-F471 cells following treatment with increasing concentrations of MRTX1133. *n* samples in each group of LKR13 cells: - = 7; 1 = 7; 10 = 3; 40 = 4; 100 = 3. *n* samples in each group of MDA-F471 cells: - = 8; 1 = 8; 10 = 7; 40 = 11; 100 = 6. n.s: non-significant (*P* > 0.05). Error- bars: standard deviations of means. *P* - values were calculated using an ordinary one-way ANOVA with Dunnett’s post-test. Results are representative of two independent experiments. **c**, Western blot analysis for the indicated proteins and phosphorylated proteins at 3 hours post- treatment to EGF without or with increasing concentrations of the KRASG12D inhibitor MRTX1133 (from Mirati Therapeutics, Inc.). Proteins were run on additional gels (4 per cell line) to separately blot with antibodies against phosphorylated and total forms of each of the indicated proteins (**Supplementary Fig. 9**). Vinculin protein levels were analysed as loading control for each gel whereby four LKR13 and four MDA-F471 blots are shown in **Supplementary Fig. 9**. For lysates from each of the two cell lines, vinculin blots from Gel 1 (**Supplementary Fig. 9**) are selected and shown in this figure panel. Uncropped images of western blots with molecular weight ladder are also shown in **Supplementary Fig. 9**. Results are representative of three independent experiments. EGF: epidermal growth factor. **d,** Size quantification of organoids derived from GFP^+^ lungs cells of NNK-treated AT2 reporter mice and treated with 200 nM MRTX1133 or control DMSO *in vitr*o (n = 6 wells per condition). Box-and-whisker definitions are similar to **Extended Data Fig. 1f**. *n* samples (organoids) in each group: DMSO = 38; MRTX1133 = 53. *P* - value was calculated using two-sided Wilcoxon Rank-Sum test. **e**, IF analysis showing representative organoids derived from sorted GFP^+^ cells from AT2 reporter mice that were exposed to saline (top two rows; n = 4 wells) or exposed to NNK and then treated *ex vivo* with DMSO (middle two rows; n = 6 wells) or 200 nM MRTX1133 (bottom two rows; n = 6 wells). Scale bars = 50 μm except for the first DMSO-treated organoid (third row) whereby scale bar = 100 μm. Staining was repeated three times with similar results.

**Extended Data Fig. 13. Analysis of labelled Krt8^+^ cells following tobacco carcinogen exposure**. **a,** Representative images of IF analysis of tdT, LAMP3, and NKX2-1 in lung tissues of control saline-treated mice (upper row; n = 2), in non-tumour (normal) lung regions of mice at end of an 8-week NNK exposure (middle row; n = 3), as well as in non-tumour (normal) lung regions of mice at 8-12 weeks following EOE to NNK (lower row; n = 3), and in *Gprc5a^-/-^*; Krt8- CreER; *Rosa^tdT^*^/+^ mice. IF analysis of tdT and Lamp3 in tumours detected in *Gprc5a^-/-^*; Krt8- CreER; *Rosa^tdT^*^/+^ mice and showing strong (**b**, n = 10) and negative/low (**c**, n = 7) tdT labelling in tumour cells. Scale bars = 10 μm.

## Supplementary Information

The **Supplementary Figures** file contains nine supplementary figures along with their legends. **Supplementary Fig. 1 outlines gating strategy for sorting epithelial cells from human lung tissues**. **Supplementary Fig. 2, 3,** and **5** include additional details pertaining to scRNA-seq data processing and analysis. **Supplementary Fig. 4 and 6 outline gating strategies for sorting lung cells from mouse models including those with AT2 lineage tracing**. **Supplementary Fig. 7** and **8** show complementary analyses in AT2 lineage-labelling mouse experiments. **Supplementary Fig. 9 shows raw western blots images for each cell line and gel analysed**. We also provide **Supplementary Tables 1-12** along with their titles.

