## Supplementary Information for "An atlas of epithelial cell states and plasticity in lung adenocarcinoma"

Departments of ^1^Genomic Medicine and ^2^Translational Molecular Pathology, ^3^Department of Pulmonary Medicine, The University of Texas MD Anderson Cancer Center, Houston, Texas. ^4^Graduate Program in Developmental Biology, Baylor College of Medicine, Houston, Texas. ^5^ The University of Texas Health Houston Graduate School of Biomedical Sciences, Houston, TX. ^6^Department of Biostatistics, Epidemiology and Informatics, Perelman School of Medicine, University of Pennsylvania, Philadelphia, PA. Departments of ^7^Thoracic, Head and Neck Medical Oncology, ^8^Cardiovascular and Thoracic Surgery, and ^9^Epidemiology, The University of Texas MD Anderson Cancer Center, Houston, TX. ^10^Department of Medicine, The University of California Los Angeles, Los Angeles, CA. ^11^Lung Cancer Initiative at Johnson and Johnson, Boston, MA. ^12^Section of Computational Biomedicine, School of Medicine, Boston University, Boston, MA.

**^*,^** ^†^ These authors contributed equally

^#^ **Correspondence**: Humam Kadara, PhD:, Linghua Wang, MD, PhD:, or Jichao Chen, PhD, MHS:.

**Supplementary Information Guide**

The Supplementary Figures file contains nine supplementary figures along with their legends. **Supplementary Fig. 1** outlines gating strategy for sorting epithelial cells from human lung tissues. **Supplementary Fig. 2, 3,** and **5** include additional details pertaining to scRNA-seq data processing and analysis. **Supplementary Fig. 4** and **6** outline gating strategies for sorting lung cells from mouse models including those with AT2 lineage tracing. **Supplementary** **Fig. 7** and **8** show complementary analyses in AT2 lineage-labelling mouse experiments. **Supplementary Fig. 9** shows raw western blots images for each cell line and gel analysed. We also provide titles description of **Supplementary Tables 1-12**.

**Outline**

Page 1 ----------------- Supplementary Fig. 1 & legend

Page 2 ----------------- Supplementary Fig. 2 & legend

Page 3 ----------------- Supplementary Fig. 3 & legend

Page 4 ----------------- Supplementary Fig. 4 & legend

Page 5 ----------------- Supplementary Fig. 5 & legend

Page 6 ----------------- Supplementary Fig. 6 & legend

Page 7 ----------------- Supplementary Fig. 7 & legend

Pages 8-9 ------------- Supplementary Fig. 8 & legend

Page 10 --------------- Supplementary Fig. 9 & legend

Page 11 --------------- List of Supplementary Tables
