## Supplementary Fig. for "An atlas of epithelial cell states and plasticity in lung adenocarcinoma"

**Supplementary Figure 1**

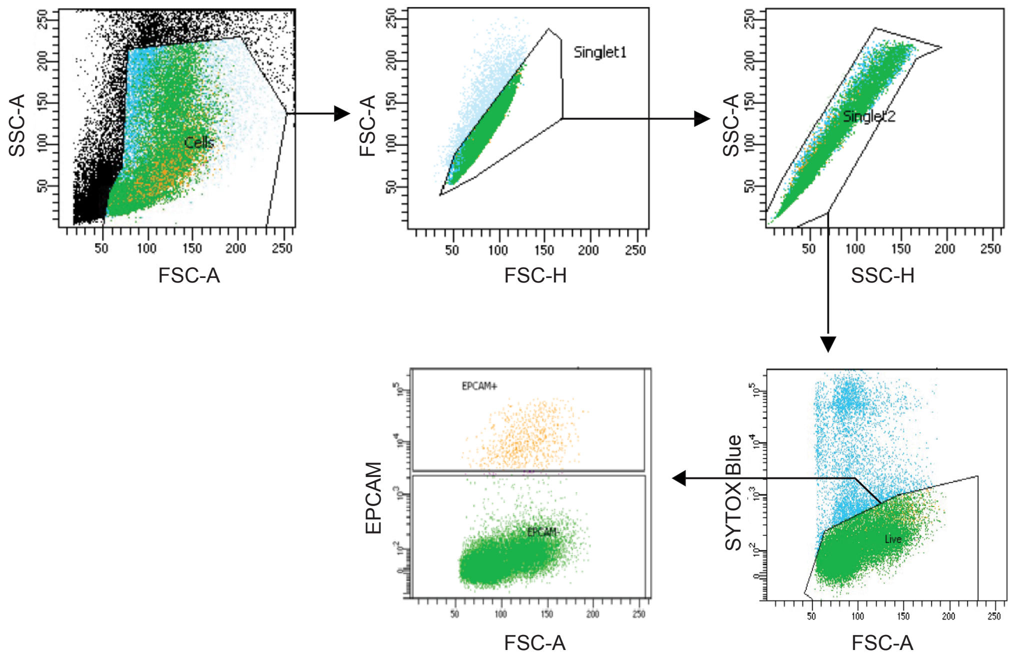


**Supplementary Fig. 1.** Gating strategy used for cell sorting of live epithelial single cells (SYTOX Blue$^{neg}$; EPCAM$^{+}$) from human normal or LUAD tissues for scRNA-seq.

**Supplementary Figure 2**

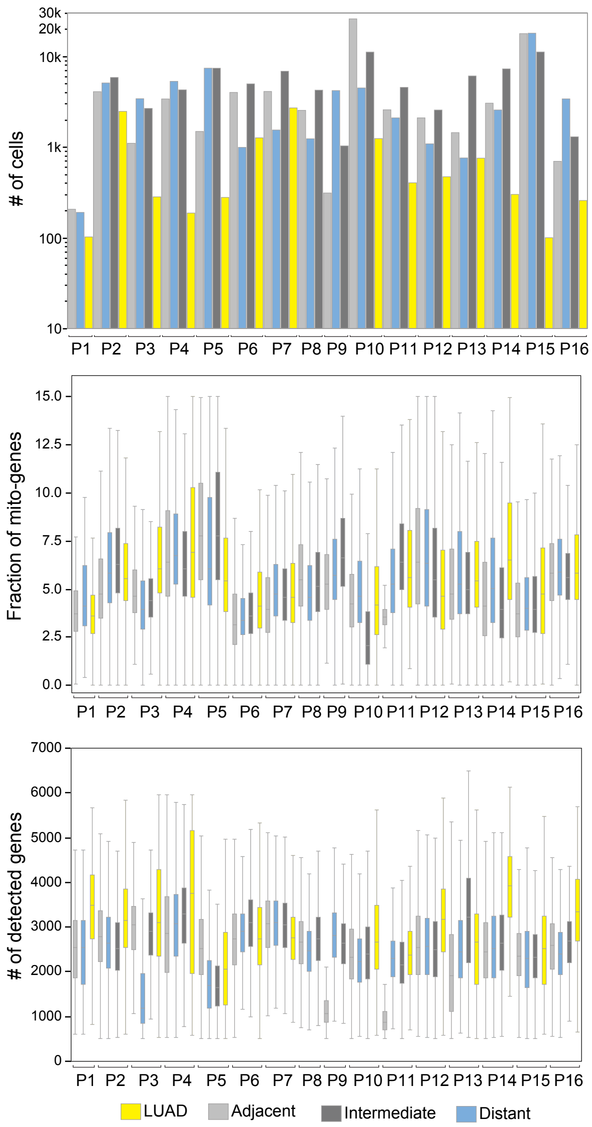


**Supplementary Fig. 2.** Quality control metrics including the number of cells retained (top), the fraction of mitochondrial genes (middle) and the number of detected genes per cell (bottom) for each sample. Y-axis is log-scaled in the top panel. Box, median ± interquartile range; whiskers, 1.5× interquartile range; centre line: median. *n* cells in each group: P1, LUAD = 166, Adjacent = 206, Distant = 190; P2, LUAD = 2,999, Adjacent = 4,071, Intermediate = 5,839, Distant = 5,076; P3, LUAD = 2,701, Adjacent = 1,099, Intermediate = 2,661, Distant = 3,407; P4, LUAD = 591, Adjacent = 3,383, Intermediate = 4,263, Distant = 5,290; P5, LUAD = 6,150, Adjacent = 1,483, Intermediate = 7,388, Distant = 7,392; P6, LUAD =3,866, Adjacent = 3,993, Intermediate = 4,962, Distant = 990; P7, LUAD = 3,518, Adjacent = 4,093, Intermediate = 6,840, Distant = 1,539; P8, LUAD = 423, Adjacent = 2,534, Intermediate = 4,234, Distant = 1,232; P9, LUAD = 549, Adjacent = 311, Intermediate = 1,028, Distant = 1,232; P10, LUAD = 1,564, Adjacent = 25,835, Intermediate = 11,127, Distant = 4,474; P11, LUAD = 778, Adjacent = 2,571, Intermediate = 4,540, Distant = 2,094; P12, LUAD = 470, Adjacent = 2,095, Intermediate = 2,556, Distant = 1,083; P13, LUAD = 793, Adjacent = 1,440, Intermediate = 6,065, Distant = 757; P14, LUAD =1,914, Adjacent = 3,041, Intermediate = 7,259, Distant = 2,558; P15, LUAD = 1,919, Adjacent = 17,769, Intermediate = 11,147, Distant = 17,962; P16, LUAD = 257, Adjacent = 695, Intermediate = 1,295, Distant = 3,390.

### Supplementary Figure 3

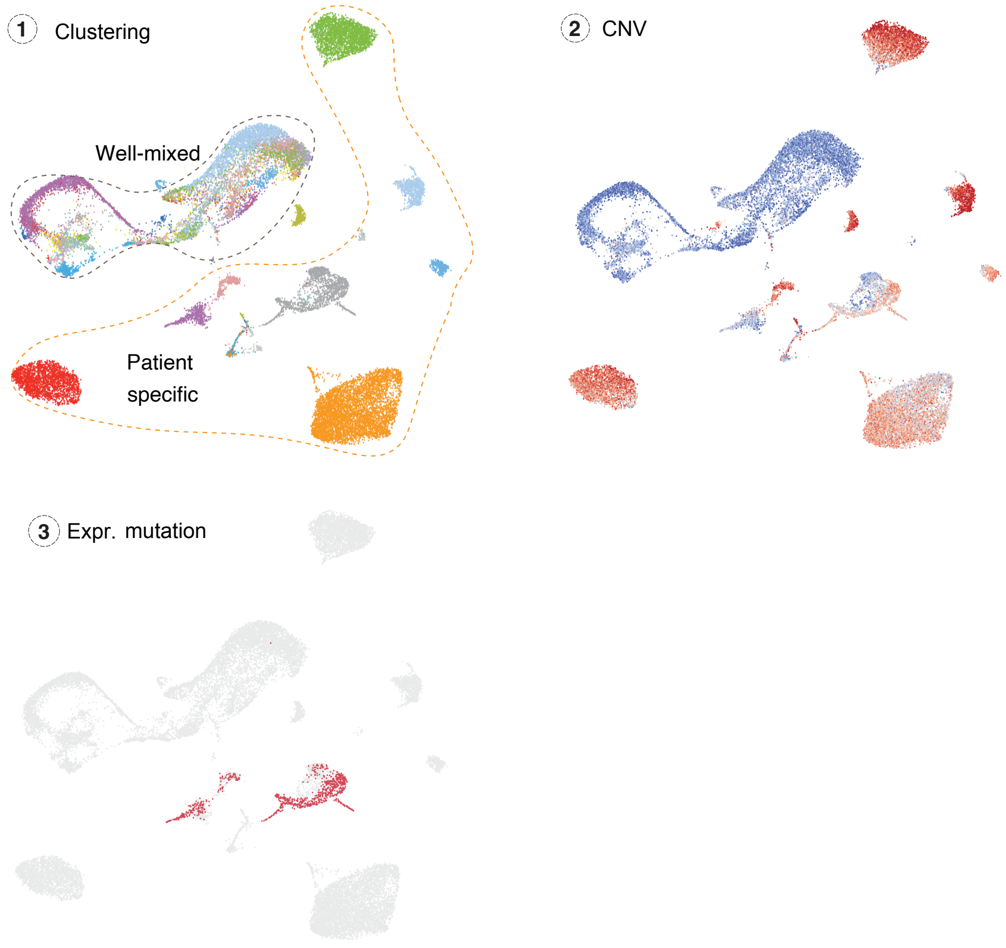


**Supplementary Fig. 3.** Schematic plot illustrating the approach used to identify malignant cells in LUAD samples based on clustering, CNV scores, and presence of *KRAS*$^{G12D}$ mutations.

**Supplementary Figure 4**

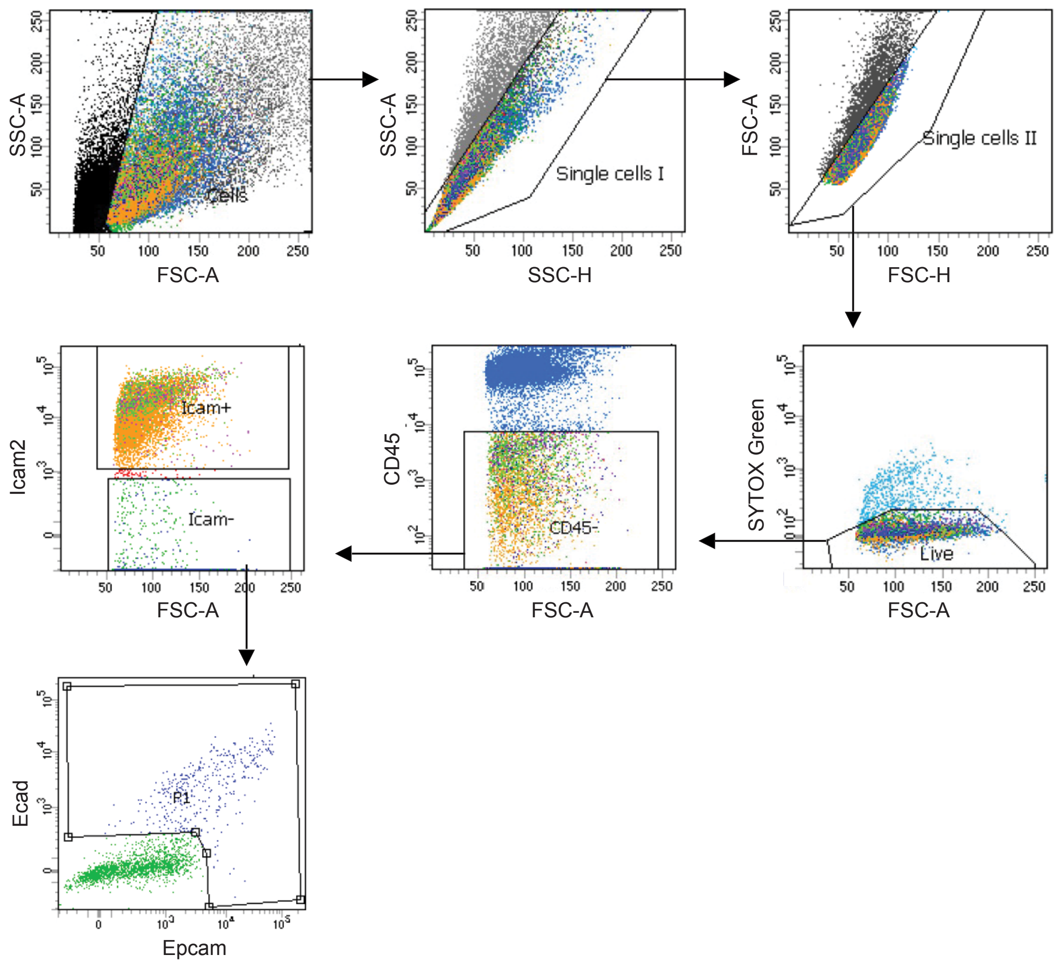


**Supplementary Fig. 4**. Gating strategy used for sorting of live epithelial single cells (SYTOX Green$^{neg}$; CD45$^{neg}$; ICAM2$^{neg}$; EPCAM$^{+}$ and/or ECAD$^{+}$) from lungs of *Gprc5a*$^{-/-}$ mice for scRNA-seq shown in **Fig. 3** and **Extended Data Fig. 7 and 10.**

**Supplementary Figure 5**

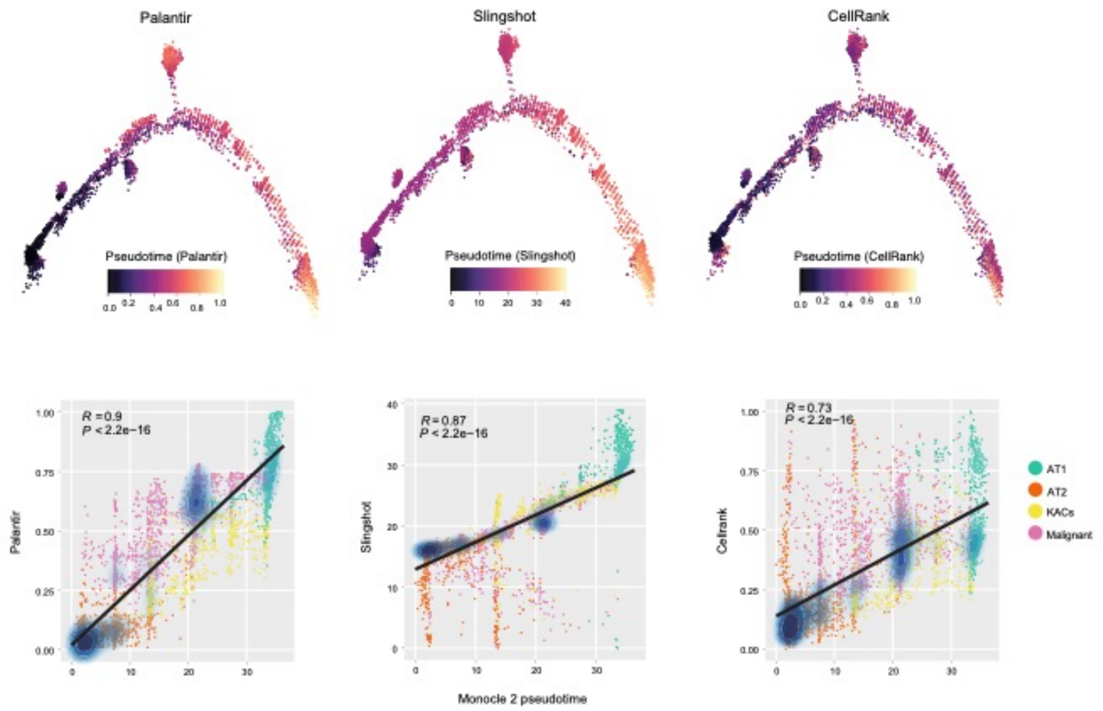


**Supplementary Fig. 5.** Robustness of the pseudotime trajectory analysis of mouse alveolar cell subsets and tumour cells. Trajectory analysis of alveolar and malignant subsets using three trajectory inference methods (top left to right: Palantir, Slingshot, and CellRank), as well as quantified correlation of each of these methods with results obtained from Monocle 2-derived pseudotime (bottom). *P* values were calculated with Spearman correlation test. R denotes Spearman correlation coefficient. The cell density estimation (see Methods) denotes the distribution density on the scatterplot.

### Supplementary Figure 6

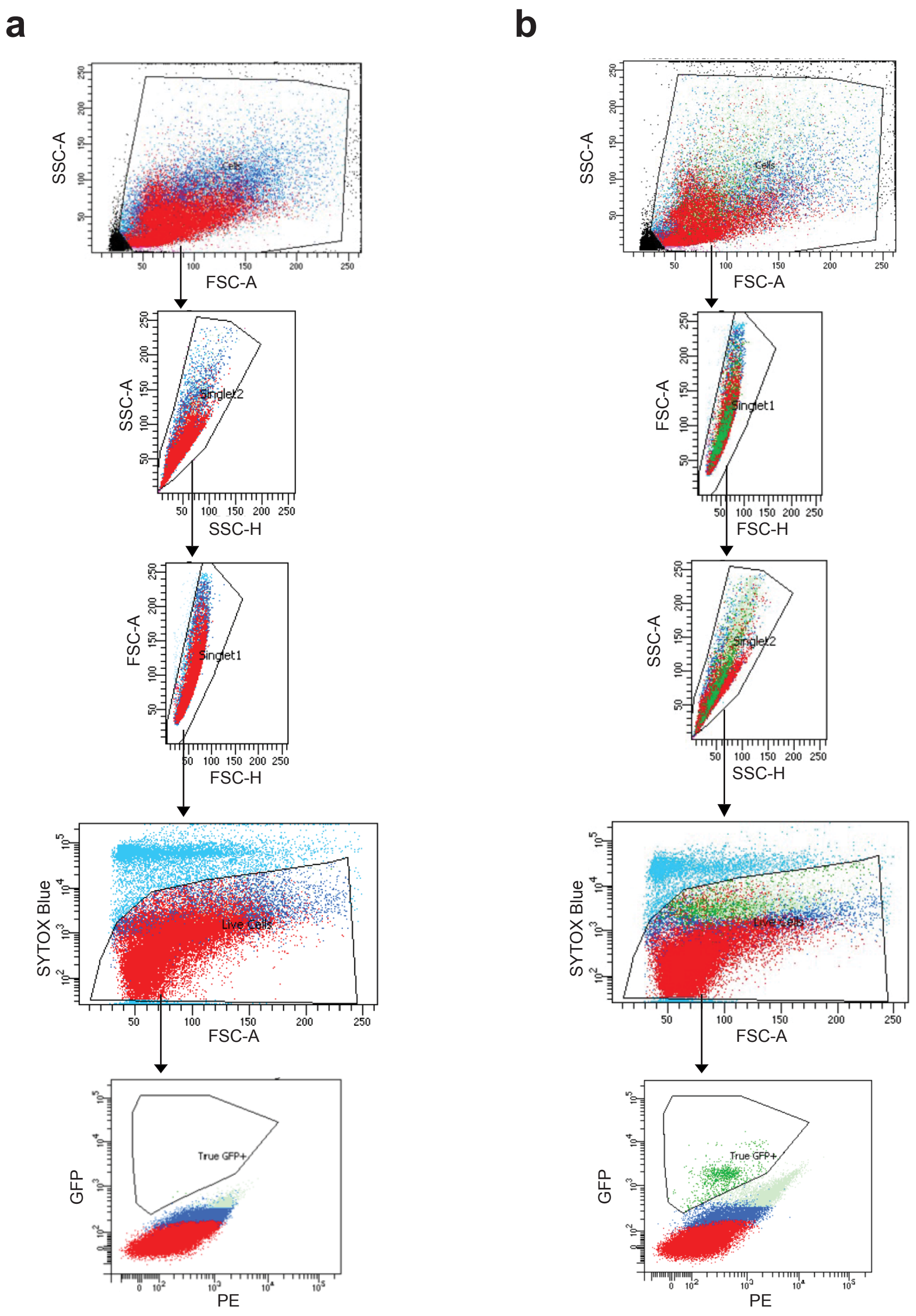


**Supplementary Fig. 6.** Gating strategy used for sorting of live GFP$^{+}$ epithelial single cells from AT2 lineage reporter mice (*Gprc5a*$^{-/-}$;*Sftpc*$^{creER/+}$;*Rosa*$^{Sun1GFP/+}$). **a**, Gating of lung cells from a *Gprc5a*$^{-/-}$;*Sftpc*$^{creER/+}$;*Rosa*$^{Sun1GFP/+}$ mouse without prior exposure to tamoxifen. This mouse was used to identify and exclude background GFP fluorescence signal from the gates set for sorting of true GFP-expressing AT2 cells in tamoxifen-exposed animals as shown in **b**. **b**, Gating strategy of lung cells from a representative tamoxifen-exposed *Gprc5a*$^{-/-}$;*Sftpc*$^{creER/+}$;*Rosa*$^{Sun1GFP/+}$ mouse to isolate live GFP-expressing cells (SYTOX Blue$^{neg}$; GFP$^{+}$) for scRNA-seq and organoid derivation shown in **Fig. 4, Extended Data Fig. 11 and 12, and Supplementary Fig. 7**.

**Supplementary Figure 7**

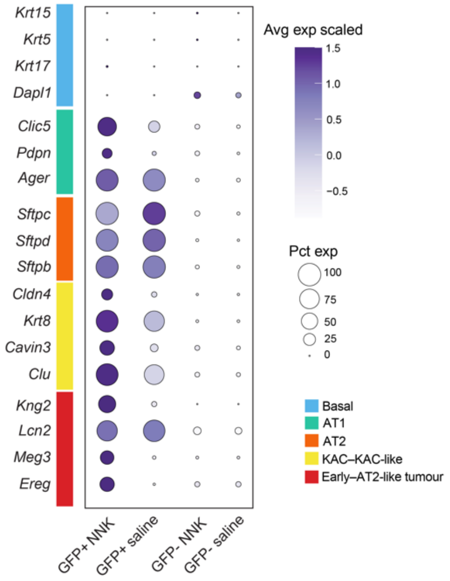


**Supplementary Fig. 7.** Proportions and average expression levels of select alveolar, tumour, and basal cell marker genes in GFP$^{+}$ and GFP$^{-}$ samples from saline- and NNK-treated AT2 lineage-labelled mice.

**Supplementary Figure 8**

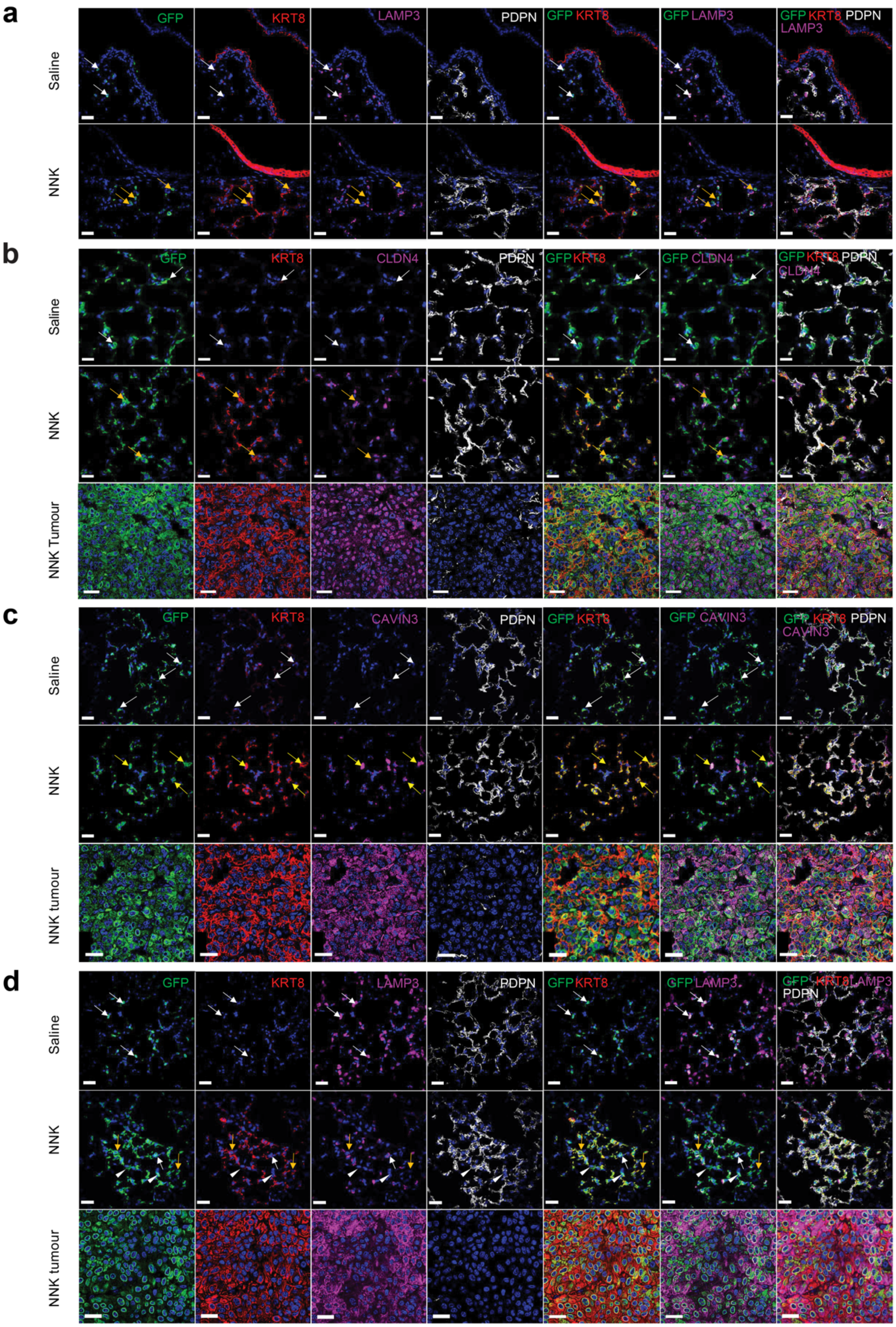

**Supplementary Fig. 8. *In situ* analysis of AT2 cells and KACs in tissues from AT2 lineage labelled tobacco carcinogenesis mouse model. a,** Representative images of IF analysis of GFP, KRT8, LAMP3, and PDPN lung parenchyma and airway epithelia in lung tissues of AT2 reporter mouse lungs at 3 months post-exposure to saline or NNK (n = 2 mice per condition). White arrows; $GFP^+$ AT2 cells. Yellow arrows; $GFP^+$ $KRT8^+$ cells. Scale bars: 30 μm. **b**, Representative images of IF analysis of GFP, KRT8, CLDN4, and PDPN in lung parenchyma and tumours of the same mice shown in **a**. White arrows; AT2 cells. Yellow arrows; KACs. Scale bars: 30 μm in first 2 rows (normal lung regions), 10 μm in the third row (tumour). **c,** Representative images of IF analysis of GFP, KRT8, the KAC marker CAVIN3, and PDPN in lung parenchyma and tumours of the same mice shown in **a**. White arrows; $GFP^+$ $KRT8^{-/low}$ AT2 cells. Yellow arrows; $GFP^+$ $KRT8^+$ KAC $marker^+$ $(CAVIN3)^+$ cells. Scale bars: 30 μm for the first two rows and 10 μm for the higher magnified tumour (third row). **d**, Representative images of IF analysis of GFP, KRT8, LAMP3, and PDPN in lung parenchyma and tumours of the same mice shown in **a**. White arrows; $GFP^+$ $KRT8^{-/low}$ AT2 cells. Yellow arrows; $GFP^+$ $KRT8^+$ cells. White arrow heads; $GFP^+$ $PDPN^+$ AT1 cells. Scale bars: 30 μm for the first two rows and 10 μm for the higher magnified tumour. Staining was repeated three times with similar results.

**Supplementary Figure 9**

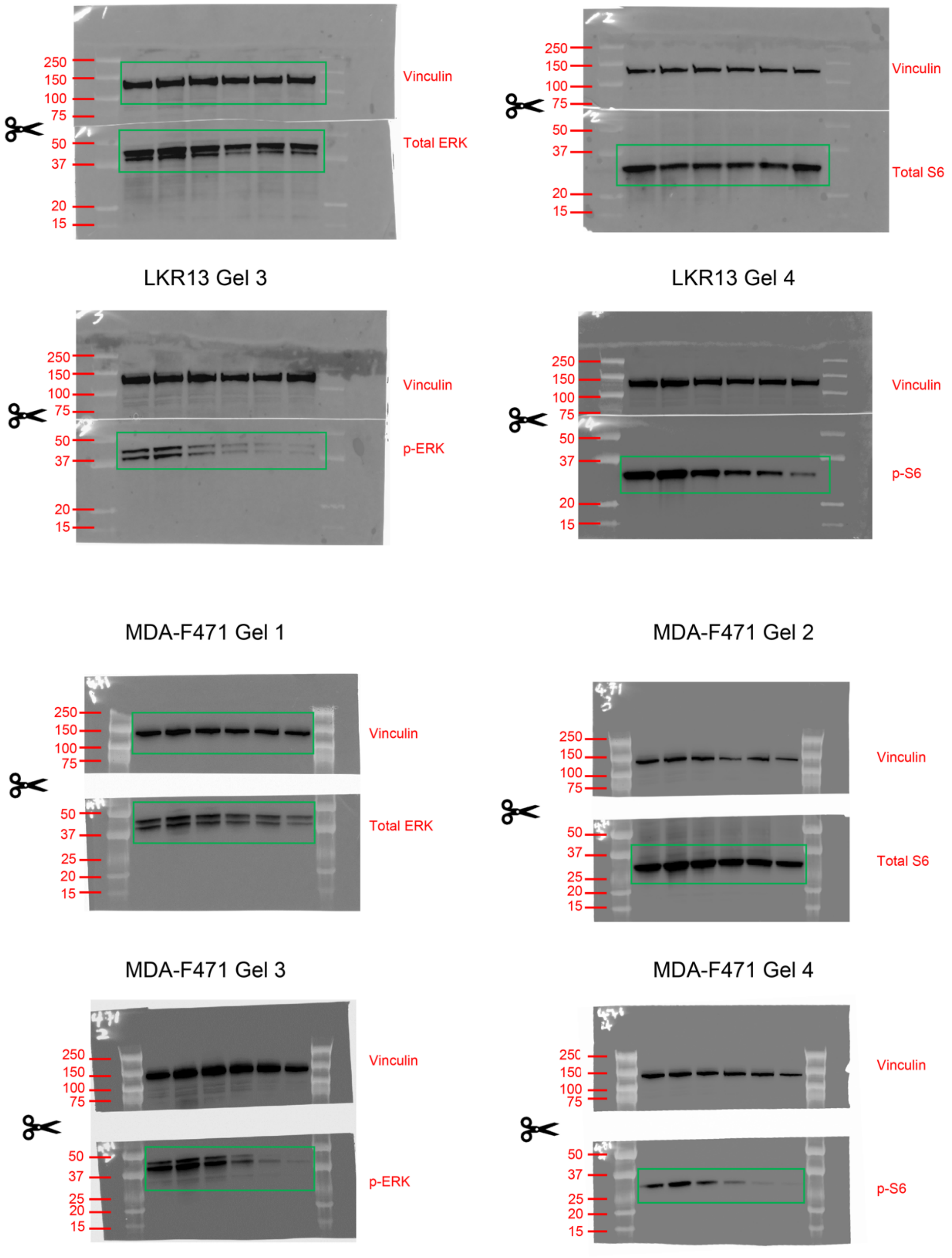


**Supplementary Fig. 9.** Western blot source data used for protein expression analysis shown in **Extended Data Fig. 12c** for LKR13 and MDA-F471 cells treated with increasing concentrations of MRTX1133 for 3 hours. Four gels were run for each set of lysates (each cell line) to separately probe for phosphorylated and total forms of S6 and ERK proteins. Vinculin was probed as loading control for each gel and only one anti-vinculin blot per cell line is shown in **Extended Data Fig. 12c.** Membranes were cut horizontally using molecular weight marker as a guide, and cut membranes were incubated with the specified antibodies. Raw images of uncropped membranes were acquired using an overlay of Chemiluminescent and Colorimetric functions obtained on the Chemidoc Touch Imaging System (Bio-Rad). Green rectangles denote blots that were included in **Extended Data Fig. 12c**.

**Supplementary Tables**

**Supplementary Table 1.** Clinicopathological features of patients with LUAD in this study.

**Supplementary Table 2.** Sample level summary of sequencing data quality metrics.

**Supplementary Table 3.** Top 50 DEGs of major epithelial lineages identified among 246,102 cells. pct.1; expression percentage of the gene in the corresponding cell lineage. pct.2; expression percentage of the gene in the other cell lineages.

**Supplementary Table 4.** Patient level summary statistics of CytoTRACE scores in 17,064 malignant cells.

**Supplementary Table 5.** Summary of human cell fractions with different driver mutations in meta-groups.

**Supplementary Table 6.** KRAS signature genes derived in this study. pct.1; expression percentage of the gene in the cell cluster (C5, **Extended Data Fig. 1f**) enriched for cells from KM-LUADs. pct.2; expression percentage of the gene in the other cell clusters.

**Supplementary Table 7.** Differentially expressed genes between human KACs and other AICs. Average log fold change and adjusted *P* – values are indicated for each gene.

**Supplementary Table 8.** Differentially expressed genes between human KACs and other alveolar cells. Average log fold change and adjusted *p* – values are indicated for each gene. pct.1; expression percentage of the gene in the KACs. pct.2; expression percentage of the gene in the other alveolar cells.

**Supplementary Table 9.** Fractions of human alveolar and malignant cells with *KRAS*$^{G12D}$ mutations.

**Supplementary Table 10.** Top 50 DEGs between lung epithelial lineages in *Gprc5a*$^{-/-}$ mice. pct.1; expression percentage of the gene in the corresponding cell lineage. pct.2; expression percentage of the gene in the other cell lineages.

**Supplementary Table 11.** Fractions of alveolar and malignant cells with *Kras*$^{G12D}$ mutations in lungs of *Gprc5a*$^{-/-}$ mice exposed to NNK or saline.

**Supplementary Table 12.** Summary statistics of inferred pseudotime scores in alveolar and tumour cells in lungs of *Gprc5a*$^{-/-}$ mice exposed to NNK or saline.
